# Design and Synthesis of Pleated DNA Origami Nanotubes with Adjustable Diameters

**DOI:** 10.1101/534792

**Authors:** Jonathan F. Berengut, Juanfang Ruan, Akihiro Kawamoto, Lawrence K. Lee

**Author notes:** Present Address: Lawrence K. Lee, EMBL Australia Node for Single Molecule Science, School of Medical Sciences, UNSW Sydney, NSW, 2052, Australia.

## Abstract

DNA origami allows for the synthesis of nanoscale structures and machines with nanometre precision and high yields. Tubular DNA origami nanostructures are particularly useful because their geometry facilitates a variety of applications including nanoparticle encapsulation, the construction of artificial membrane pores and as structural scaffolds that can spatially arrange nanoparticles in circular, linear and helical arrays. Here we report a simple computational approach that determines minimally-strained DNA staple crossover locations for arbitrary nanotube internal angles. We apply the method in the design and synthesis of radially symmetric DNA origami nanotubes with arbitrary diameters and DNA helix stoichiometries. These include regular nanotubes where the wall of the structure is composed of a single layer of DNA helices, as well as those with a thicker pleated wall structure that have a greater rigidity and allow for continuously adjustable diameters and distances between parallel helices. We also introduce a DNA origami staple strand routing that incorporates both antiparallel and parallel crossovers and demonstrate its application to further rigidify pleated DNA nanotubes.

## INTRODUCTION

DNA nanotechnology utilizes the well-known structural properties and complementary base-pairing rules of DNA (1) for the self-assembly of rationally designed nanoscale structures and machines (2–8). DNA strands at specific sites on these structures can be functionalized to selectively bind to small molecules such as nanoparticles, dyes and proteins to control their spatial organization at resolutions well below 10 nm (9–11). Thus, DNA nanostructures are suitable for a broad range of applications. For example, metallic nanoparticles can be spatially arranged to construct DNA-based plasmonic architectures (12–14) for fluorescence enhancement (15) or surface enhanced Raman scattering (16–18), which can be used as highly sensitive molecular sensors (19–21). In addition, immobilization of biomolecules allows for the construction of multi-enzyme complexes (22–26), as well as the control of biomolecular assembly (27–30) and cellular processes (31–33).

Tubular DNA structures have properties that make them particularly useful (34). Their hollow structure can be used to construct biomimetic membrane channels (30,35–37), to encapsulate proteins for multi-enzyme bioreactors (24,25) or to selectively deliver cargo to, and mediate the activity of specific cell types (32,38). DNA nanotubes have greater structural rigidity than single DNA duplexes (39). This makes them suitable for such applications as the alignment of proteins in solution for nuclear magnetic resonance spectroscopy (40), the construction of molecular barcodes for calibration of super-resolution microscopy methods (41,42), or for scaffolding extended linear arrays of metallic nanoparticles (43–45), which is useful for the bottom-up construction of nanowires (46,47). In addition, nanotubes can organize nanoparticles into circular and helical arrays, which can be used to construct plasmonic nanostructures (48,49) with distinct optical resonances that depend on their chirality (12).

DNA nanotubes with defined diameters can be synthesized from repeating arrays of short DNA motifs (50–53). The length of these nanotubes cannot be controlled however, and each site on the nanotube is not uniquely addressable. To create nanotubes with fixed dimensions and full addressability, the DNA origami method can be used (4). This involves folding a long single-stranded DNA ‘template’ into a desired structure by hybridization to many shorter ‘staple’ strands, forming arrays of double-stranded DNA helices linked by crossovers, which are junctions in which staple or template strands cross from one helix to another (Figure 1A). A canonical DNA origami rectangle consists of a parallel array of DNA helices through which the template strand weaves back and forth in opposite directions on adjacent helices (Figure 1B). This means that structures tend to consist of an even number of helices to avoid long sections of unpaired bases. Nanotubes can typically be constructed in one of two configurations; by bending the DNA helices and arranging them in coaxial or helical arrays (6,54,55), or by incorporating additional staple crossovers linking the first and the last helix of a rectangle to arrange them in parallel arrays (48,49). The latter configuration can be thought of as the ‘rolling up’ of a DNA origami rectangle along an axis parallel to the DNA helices, forming a hollow cylinder with walls the thickness of a single layer of DNA helices (Figure 1C). As with the rectangle, each adjacent helix is antiparallel and the nanotube has an even number of helices. If the staples and crossovers that connect the rectangle’s helices together are arranged in a repeating array, the resulting nanotube will have radial symmetry. This configuration allows for a discreet set of possible nanotube diameters that are dependent on the number of helices in the DNA origami rectangle.

**Figure 1.**
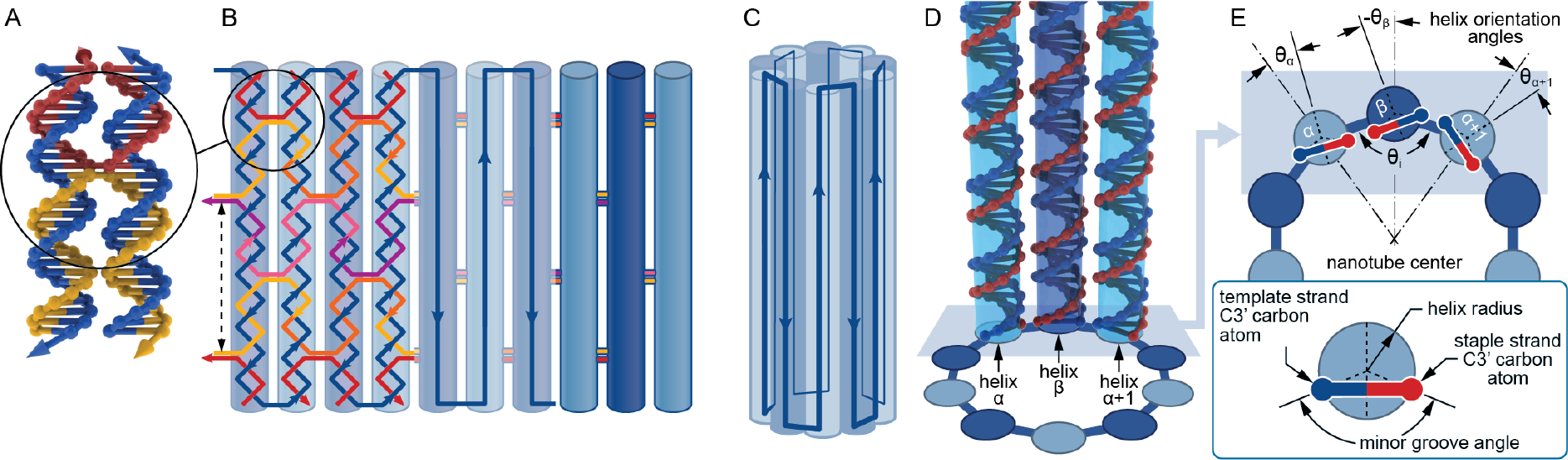
Schematic depiction of staple routing and architecture of radially symmetric DNA origami nanotubes. (A) A section of two adjacent DNA helices joined by reciprocal staple crossovers. The direction of the DNA strands in the 3’ to 5’ direction is indicated by an arrow at the ends of the strands. (B) An unrolled DNA origami nanotube showing the S-shaped staple routing as well as the direction in which the template strand weaves through the nanotube. Helices are shown in light and dark blue according to the template strand direction. Dashed arrow indicates the ‘length unit’ that would be repeated along the length of a longer nanotube. (C) An assembled DNA origami nanotube. (D,E) Position and rotational orientation of helices α, β, and α+1. The helix orientation angles θ_α_, θ_β_ and θ_α+1_ and the internal angle θ_i_ are indicated. Since helices α and α+1 are radially symmetric about the nanotube center, θ_α_ is equal to θ_α+1_.

To define the exterior and interior surfaces of the nanotube, it is useful to rationally control the internal angle between adjacent helices (49) (Figure 1D). This can be done by altering the periodicity of these crossovers, according to:

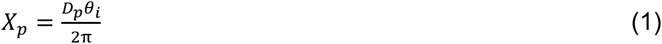

where X_p_ is the crossover periodicity, θ_i_ is the internal angle between any three sequential helices and D_p_ is the periodicity of a DNA duplex, approximately 10.5 base-pairs (bp) per turn (4,56,57). For a regular DNA nanotube with *n* helices, θ_i_ is the internal angle of a regular polygon with *n* sides:

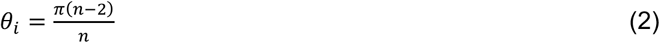

and hence by substitution, the crossover periodicity for a DNA nanotube with *n* helices can be defined as (58):

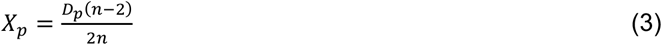

For example, a hexagonal nanotube with an internal angle of 120° has a crossover periodicity of 3.5, which can be achieved by locating staple crossovers every 7 bp or some multiple thereof (5). Since the ends of staple crossovers cannot physically occur in between base pairs, crossovers need to be positioned at an integer multiple of the crossover periodicity. However, in practice, there are few *n* for which the desired crossover periodicity is a multiple of a low integer number of base pairs (58). This is required to ensure sufficient crossovers to construct a well-formed nanotube. To create arbitrary *n*-helix nanotubes with minimal strain, crossovers can be incorporated at multiple periodicities, where the global internal angle is a consequence of the average of all crossover periodicities (49). This inevitably requires that some crossovers will occur at locations where the unstrained location of proximal DNA bases on adjacent helices do not align with the geometry of a crossover. Subsequently, the DNA must twist or bend to accommodate the crossover resulting in a strained structure. Determining the crossover locations that achieve both the desired periodicity and minimal strain is a key challenge for DNA nanostructure design (6). The solution depends not only on the desired internal angle, but also the relative orientation around and translation along the axis of each individual helix.

Here we address the challenge of designing DNA origami nanotubes with an empirical approach that readily identifies minimally-strained crossover locations for nanotubes with arbitrary internal angles. The approach is applied to the design and synthesis of a regular rolled-rectangle nanotube, and for DNA origami nanotubes with a pleated wall structure similar to those conceptualized theoretically (58). The latter design allows for nanotubes with an arbitrary and continuous set of diameters and interhelical spacing, and multi-layer DNA origami walls providing additional rigidity over single-layer walls, which lack structural rigidity (57). Finally, we present a new staple routing that adds further rigidity, provides additional control over the dimensions of the pleated nanotube and which is generally applicable to other DNA nanostructures.

## MATERIAL AND METHODS

### Calculation of predicted nanotube geometry

The predicted outer diameters of the nanotubes were determined by:

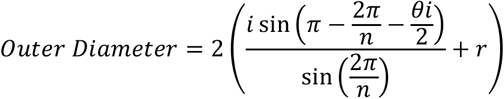

Where *n* = number of helices in nanotube, *i* = interhelical distance (set to 3 nm), θ_i_ = internal angle, and *r* = DNA double helix radius (set to 1 nm). The predicted length of a DNA nanotube was calculated by multiplying the number of base pairs in the longest DNA helix by the rise between DNA bases along the DNA helix 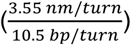.

### DNA origami synthesis

To prepare DNA origami samples for AFM, 5 nM M13mp18 ssDNA template strands (Bayou Biolabs) was mixed with 5-fold molar excess of DNA staple strands (Integrated DNA Technologies) in DNA origami synthesis buffer 1 (33 mM Tris Acetate, 12.5 mM Mg Acetate, pH 8.2) and annealed with temperature ramp 1 (95 °C for 10 mins, then 94 to 86 °C over 10 mins in 0.1 °C steps, 85 to 70 °C over 75 mins in 0.1 °C steps, 70 to 40 °C in 450 mins in 0.1 °C steps, 40 to 25 °C in 150 mins in 0.1 °C steps, then 10 mins at 20 °C and stored at 4 °C until use). To prepare DNA origami samples for imaging with electron microscopy 20 nM M13mp18 ssDNA template strands was mixed with 10-fold molar excess of DNA staple strands in DNA origami buffer 2 (5 mM Tris, 1 mM EDTA, 16 mM MgCl2, pH 8.0) and annealed with temperature ramp 2 (65 °C for 15 mins, then 60 to 40 °C in 200 steps at 5 mins/step, then 40 to 25 °C in 150 steps at 1 min/step).

### Atomic force microscopy

DNA origami samples were either purified with size exclusion chromatography in hand-packed Micro Bio-Spin™ Columns (Bio-Rad) loaded with 750 μl of Sephacryl™ S-300 HR beads (GE Healthcare), which had an exclusion limit of 300 kDa, or purified using PEG precipitation (60). Purified samples were diluted between 1:2 and 1:5 in DNA origami buffer 1. A drop of sample (5 μl) was placed onto freshly cleaved mica surface and allowed to adsorb for 5 minutes, then AFM head was lowered until contact between the tip and the sample in solution was made. Imaging was performed on a Bioscope Catalyst™ Atomic Force Microscope (Bruker) using ScanAsyst and Peakforce Tapping mode in fluid, with SNL-A tips (Bruker). Spring constant was set to between 0.35 N/m & 0.58 N/m, scan rate was between 0.5 Hz & 1.0 Hz. Scan size was typically 1 μm square with 256 or 512 samples/line. Images were processed using Gwyddion (61) by first levelling data by mean plane subtraction, then aligning rows by matching height median, then setting the zero height and levelling again by fitting a plane through three points that did not contain material to define the background intensity. Each of the three points was chosen in an area free from structures and averaged over an 11-pixel radius. Measurements were made manually using the measure tool. Histograms were created in Wolfram Mathematica version 11.2.

### Transmission electron microscopy

DNA origami nanotubes were purified using PEG precipitation (60) and drop cast onto carbon/formvar coated copper grids and given 4 minutes to adsorb. Excess sample was wicked off using filter paper. A drop of 2% aqueous uranyl acetate was then applied to the grid and immediately wicked off using filter paper. Grids were dried overnight or until imaging. Imaging was performed on a Tecnai G2 20 Transmission Electron Microscope (FEI) in bright field mode at 200 kV. Images were analyzed in ImageJ (62). Histograms were created in Wolfram Mathematica version 11.2. Particle averages of TEM micrographs were obtained using Relion (63).

### Cryo-electron microscopy

DNA origami nanotubes were imaged with cryo-EM with one of two different conditions. In the first, 4μL of PEG-precipitation purified DNA origami were applied on glow-discharged Quantifoil R2/2 copper grids (Quantifoil Micro Tools) and plunge frozen in liquid ethane cooled liquid nitrogen using a Lecia EM GP device (Leica Microsystem). The grids were imaged using a Talos Arctica transmission electron microscope (Thermo Fisher Scientific) and operated at 200kv, with the specimen maintained at liquid nitrogen temperatures. Images were recorded at magnification 92k× on a Falcon 3EC direct detector camera operated in linear mode.

In the second condition, Quantifoil molybdenum 200 mesh R1.2/1.3 holey carbon grids were glow discharged for 20 seconds. A 3 μl aliquot of the sample solution was applied onto the grid, was blotted by filter papers for 8 sec at 100% humidity and 4 C, and the grid was quickly frozen by rapidly plunging it into liquid ethane using Vitrobot Mark IV (Thermo Fisher Scientific). The grid was inserted into a Titan Krios FEG transmission electron microscopy (Thermo Fisher Scientific) operated at 300 kV with a cryo specimen stage cooled with liquid nitrogen. Cryo-EM images were recorded with a FEI Falcon II 4k × 4k CMOS direct electron detector (Thermo Fisher Scientific) at a nominal magnification of 75k×, corresponding to an image pixel size of 1.07 Å. Images were acquired by collecting seven movie frames with a dose rate of 45 electrons per square angstrom per second and an exposure time of 2 sec.

## RESULTS AND DISCUSSION

We developed an empirical approach for the design of minimally-strained DNA origami nanotubes with an arbitrary even number *n* of helices and *n/2* radial symmetry. The approach computationally searches for optimum locations for staple strand crossovers along the length of a geometric model of the nanotube and for template strand crossovers at either end of the nanotube. For the template-strand crossovers, a strain score is calculated as the sum of the squared difference between measured and ideal distances (0.68 nm) between the nucleoside C3’ carbon atoms at crossover locations (Figure S1A). Since staple crossovers occur in pairs that reciprocate at each crossover location, distances between two successive bases on each helix must be accounted for (Figure 1A and B). The midpoint between successive C3’ carbon atoms is known as the nucleoside end midpoint (NEMid) (58) and the strain score at staple crossovers was defined as the sum of the squared difference between measured and ideal inter-NEMid distances (Figure S1B and C). The total nanotube strain score is the sum of the strain scores from all staple and template strand crossovers.

The search locates staple crossovers approximately every 1.5 turns of the DNA helix where the exact crossover periodicity is dependent on the desired internal angle (see Supplementary Note 1). This allows for staples to adopt the canonical S-shape where each staple crosses-over between three adjacent helices (Figure 1B). This design reduces the probability of poorly incorporated staples from kinetic traps (58) and ensures that nanotubes have a high density of crossovers for structural integrity. Consequently, the length of the nanotube can only be incremented in discreet ‘length units’ of approximately three turns of a DNA helix, within which each helix has two staple crossovers, one to each neighbouring helix, and two sets of staples running in opposite directions around the nanotube (Figure 1B).

The overall strain and position of staple crossovers depends on the relative rotation of each DNA helix around its helical axis. Thus, to design a nanotube with the desired even number of *n* DNA helices, the internal angle is fixed according to equation (2) and the search is carried out for all possible rotational orientations of the helices. The approach thus yields the minimally-strained crossover locations for the synthesis of radially-symmetric nanotubes with an arbitrary number of helices and a defined internal angle. Since the DNA nanotube has *n*/2 radial symmetry, all crossover locations in the structure can be found by locating the best crossovers between one DNA helix (helix β) and its two neighbours (helix α and helix α+1) (Figure 1D). This is because for any given internal angle, helix α is symmetrically equivalent to helix α+1, therefore the crossover locations between helix α and helix β also defines the crossover locations between helix α+1 and helix β+1. Similarly, to identify the minimally-strained nanotube design, it is only necessary to search the possible rotational arrangements of helix α and helix β (Figure 1E).

2D heat maps of strain scores for all θ_α_ and θ_β_ for a range of nanotube sizes from 6- to 96-helix nanotubes as well as for a flat sheet are shown in Figure S2. All structures have θ_α_ and θ_β_ angles that predict a high degree of strain as well as wide plateaus of ‘allowable’ angles where the predicted strain is relatively low. Within this plateau the set of crossover locations that result in a minimally-strained structure can be identified. Each DNA origami structure also has two continuous sets of scores each associated with a different direction around which the template strand threads through the nanotube. Sharp lines in heat maps mark the θ_α_ and θ_β_ angles at which the lowest scoring structure switches between these two template directions. For a flat sheet, scores for either template direction are equivalent because without curvature the sheet is symmetrical and template routing in either direction results in the same structure. However, with increasing internal angle, there is an increasing preference for one template strand direction over the other. This is because structures where minor grooves tend to occur closer to the inner surface of the nanotube more easily satisfy the geometric constraints of crossovers.

### Design, synthesis and characterization of regular nanotube

The search algorithm was used to design and synthesize a minimally-strained regular 20-helix nanotube with a defined inside and outside surface (Figure 2A and B). Staple crossovers occurred every 15 bp, but with every fourth crossover at 16 bp resulting in an average staple crossover periodicity of 15.25 bp, which is also similar to one turn of the DNA duplex at 10.5 bp/turn plus the ideal crossover periodicity for a 20-helix nanotube (15.23 crossovers/turn from equation 3).

**Figure 2.**
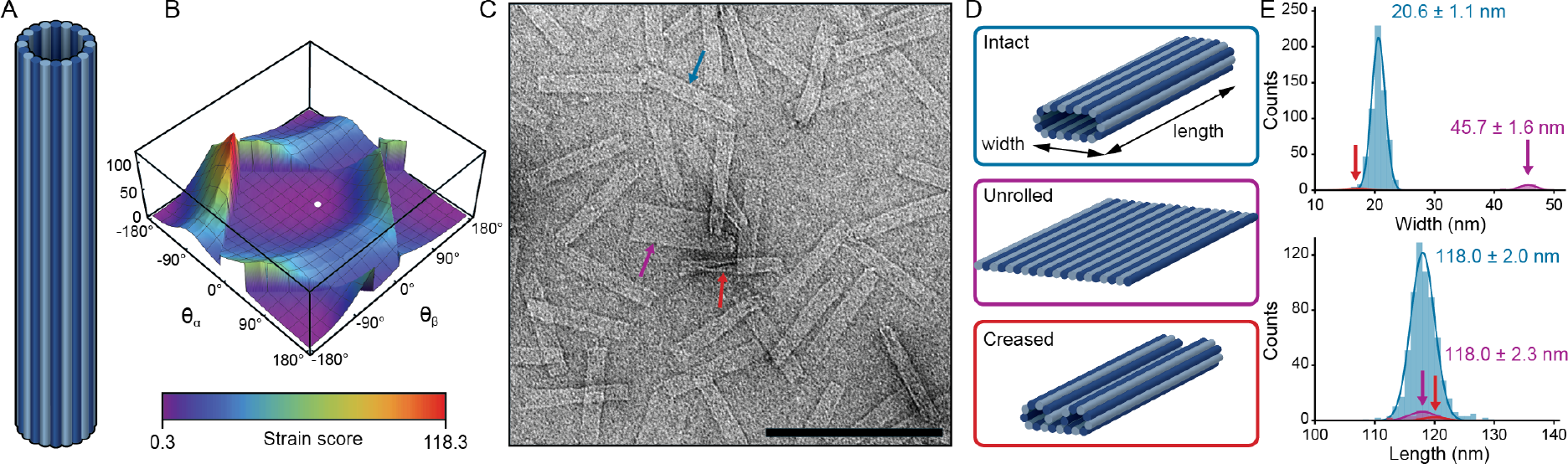
Design and synthesis of a regular 20-helix nanotube. (A) Illustration of the nanotube design. DNA helices are shown as cylinders and shaded according to the direction of the template strand. (B) 3D plot of strain scores for all θ_α_ and θ_β_ helix orientations on the nanotube. The orientation that yielded the least-strained structure is indicated by a white dot. (C) Typical TEM micrograph of the 20-helix nanotube with blue, purple and red arrows pointing to example of intact, unrolled and creased structures respectively. Scale bar represents 200 nm. (D) Models of the three categories of structure observed in TEM micrographs; intact, unrolled and creased. (E) Histograms of lengths and widths measured from TEM micrographs fitted with a Gaussian function. Data from nanotubes assessed as intact, unrolled and creased are shown in blue, purple and red respectively. Labels are average ± 1 SD. Similar measurements by AFM and cryo-EM are shown in Figure S3.

Nanotubes were characterized with transmission electron microscopy (TEM), atomic force microscopy (AFM) and cryo-electron microscopy (cryo-EM) (Figure 2C; Supplementary Figure S3). With all imaging methods, the nanotubes appeared mostly as uniform rectangular structures whose lengths (TEM = 118.1 ± 2.0 nm, cryo-EM = 118.4 ± 2.9 nm, AFM = 121.1 ± 7.6 nm) (Figure 2E; Supplementary Figure S3) were consistent with an ideal model assuming a helical pitch of 3.55 nm/turn (119.7 nm) for a DNA duplex. In TEM and cryo-EM micrographs, striations can be seen running along the length of the nanotube, confirming the direction of the DNA helices. The measured widths of the intact structures, as measured with TEM and cryo-EM (20.6 ± 1.1 and 19.4 ± 1.7 nm respectively) were similar to the diameter of an ideal model (21.2 nm), whereas widths measured with AFM were consistent with a flattened nanotube (30.4 ± 2.6 nm)(Supplementary Figure S3), which likely results from affinity to the surface and mechanical compression by the AFM tip (49,57). This interaction also likely contributed to the higher ratio of structures which appear to have split and unrolled in AFM (13.2%) as compared to those seen in TEM (5.3%) micrographs (49). A smaller proportion of structures appeared to be creased or deformed (2.2%) in TEM micrographs (Figure 2C and D). These were not seen in AFM or cryo-EM micrographs and may be associated with TEM staining.

### Design, synthesis and characterization of pleated nanotubes

One benefit of this empirical approach to locating staple and template strand crossover locations is in the ease of its use for the design of more complicated nanotube structures. For example, the internal angle can be arbitrarily set to any value that doesn’t result in helices sterically clashing. The maximum internal angle is defined by equation (2) and represents a special case for creating a regular polygonal nanotube, whereas any other internal angle should result in nanotubes where helices form a zig-zag or “pleated” wall (Figure 3A) (58). This pleated design offers several benefits over a non-pleated structure. Firstly, the distance between every second helix can be set arbitrarily from a maximum of ~6 nm in a flat sheet where θ_i_ = 180° to a minimum of ~3 nm, where θ_i_ = 60° and helices α, β and α+1 form an equilateral triangle. By altering θ_i_, nanotubes with arbitrary diameters can also be synthesized. Secondly, this pleated arrangement creates an internal ring of parallel helices where the template strand runs in the same direction and can occur in both even and odd symmetries. One possible application for these nanotubes is to construct scaffolds for the spatial arrangement of nanoparticles or biological molecules in ring-like arrays (28), and so the ability to create odd-numbered but symmetrical arrays with adjustable diameters could prove very useful. A third advantage of this pleated design is that nanotubes will have a thicker two-helix wall, which should result in increased structural rigidity. When θ_i_ = 60° a nanotube structure should be at its most rigid, as the inner helices of the nanotube will be in close proximity and hence have a more restricted range of motion.

**Figure 3.**
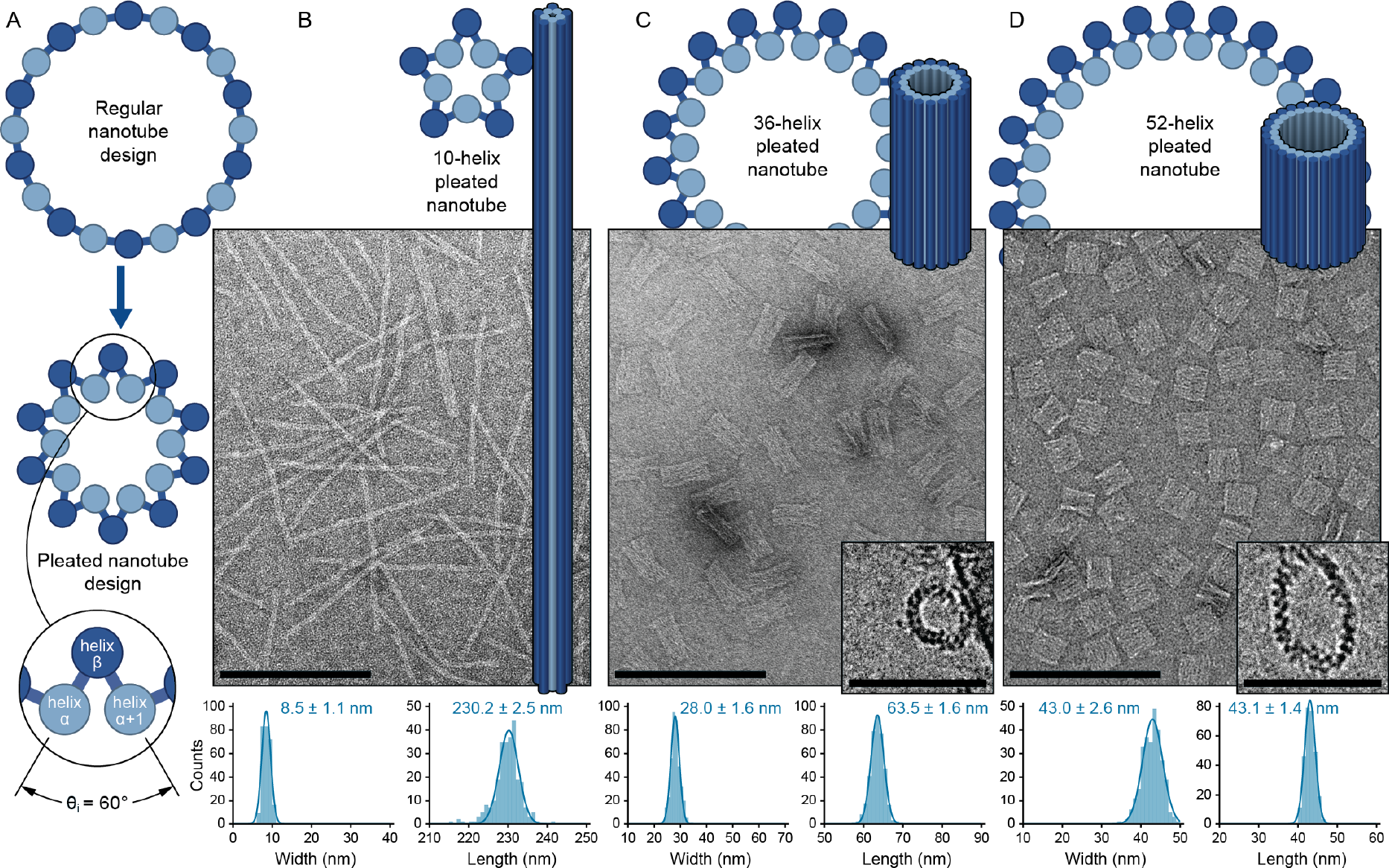
Design and synthesis of pleated nanotubes. (A) Schematic cross-sectional view of the regular (top) and pleated 20-helix nanotube (middle) with a detail view of the pleated structure showing an internal angle of 60° (bottom). Data for the 10-helix, 36-helix and 52-helix pleated nanotubes are shown in panels B, C and D respectively. Top: Schematic and pictorial views. Middle: Representative TEM images (Scale bar: 200 nm) with insets in C and D showing end-on view from a cryo-EM micrograph (Scale bar: 100 nm). Bottom: Histograms of widths and lengths of nanotubes measured with TEM with average ± 1 SD labelled.

We altered the algorithm to allow for the design of arbitrary internal angles and used the approach to design several pleated nanotubes with an internal angle of 60°. Strain score heat maps for pleated nanotubes with a range of diameters is shown in Figure S4. We synthesized minimally-strained 10-, 36- and 52-helix pleated DNA origami nanotubes. Since the same template strand was used for all designs (M13mp18, 7249 nucleotides), the lengths of the nanotubes correlated inversely with the number of helices around the circumference. The relative stability of nanotubes was assessed with AFM (Supplementary Figure S5A and B). The regular 20-helix nanotube was the most resilient to unrolling by AFM (13.2% unrolled) followed by the 36-helix (26.4%) and 52-helix (33.6%) pleated nanotubes. We attribute this to the lower number of crossover strands along the length of the shorter nanotubes. The 10-helix nanotubes tended to be physically displaced by the AFM tip, which made their shape difficult to assess (data not shown). In TEM micrographs, 10-helix pleated nanotubes appear as thin filamentous structures whose length and width corresponded to their design (Figure 3B). 36- and 52-helix pleated nanotubes appeared as rectangles with visible striations along their lengths, which matched the predicted lengths (Figure 3C and D). The 36-helix nanotubes were noticeably wider at their ends, giving the nanotubes an “hourglass” shape. This may be due to a global chirality in the nanotube caused by crossover periodicity that is not aligned with that of a DNA helix (4,49,57). Both the 36- and 52-helix pleated nanotubes were larger than their predicted diameters suggesting that pleated nanotubes are also somewhat flattened on the TEM grid (Figure 3C and D; Supplementary Table S1).

We also imaged the pleated DNA nanotubes with cryo-EM to measure their dimensions in solution. To avoid any surface effects, only nanotubes that were free from the carbon support of the cryo-EM grid were measured, although many nanotubes were associated with these supports (Supplementary Figure S6). The width of the 36-helix pleated nanotube was consistent with the predicted diameter, whereas the 52-helix pleated nanotube was on average 25% wider (Supplementary Table S1). The shorter aspect ratio of the pleated nanotubes also allowed some to be visualized end-on, with the nanotube axis perpendicular to the imaging plane. These end-on images allowed direct measurement of the nanotube’s circumference, as well as visualization of the pleated geometry (Figure 3C and D, Supplementary Figures S6 and S7) and quantification of interhelical distances (36-helix = 3.0 ± 0.3 nm, 52-helix = 3.2 ± 0.3 nm). End-on images of pleated nanotubes often deviated from a circular shape, indicating that pleated nanotubes were likely to have a degree of flexibility in solution (Figure S6). The diameter of the 36-helix nanotube calculated from circumference measurements was consistent with the measured widths of the same structures, but the measured circumferences of the 52-helix nanotubes yielded diameters smaller than the measured widths (Supplementary Table S1). This difference may be explained by the thin vitreous ice sheet in which nanotubes were frozen for cryo-EM experiments. Since the thickness of this sheet is comparable to the diameter of the 52-helix pleated nanotube, it is possible that surface effects at the air-water interface resulted in these nanotubes being compressed. Notably, the diameter of the 52-helix pleated nanotube was by all measurements substantially wider than predicted. We suggest that this is due to electrostatic repulsion between tightly-packed helices of the pleated nanotubes resulting in internal angles greater than 60° and an expansion in the nanotube diameter (Figure 4A and B).

**Figure 4.**
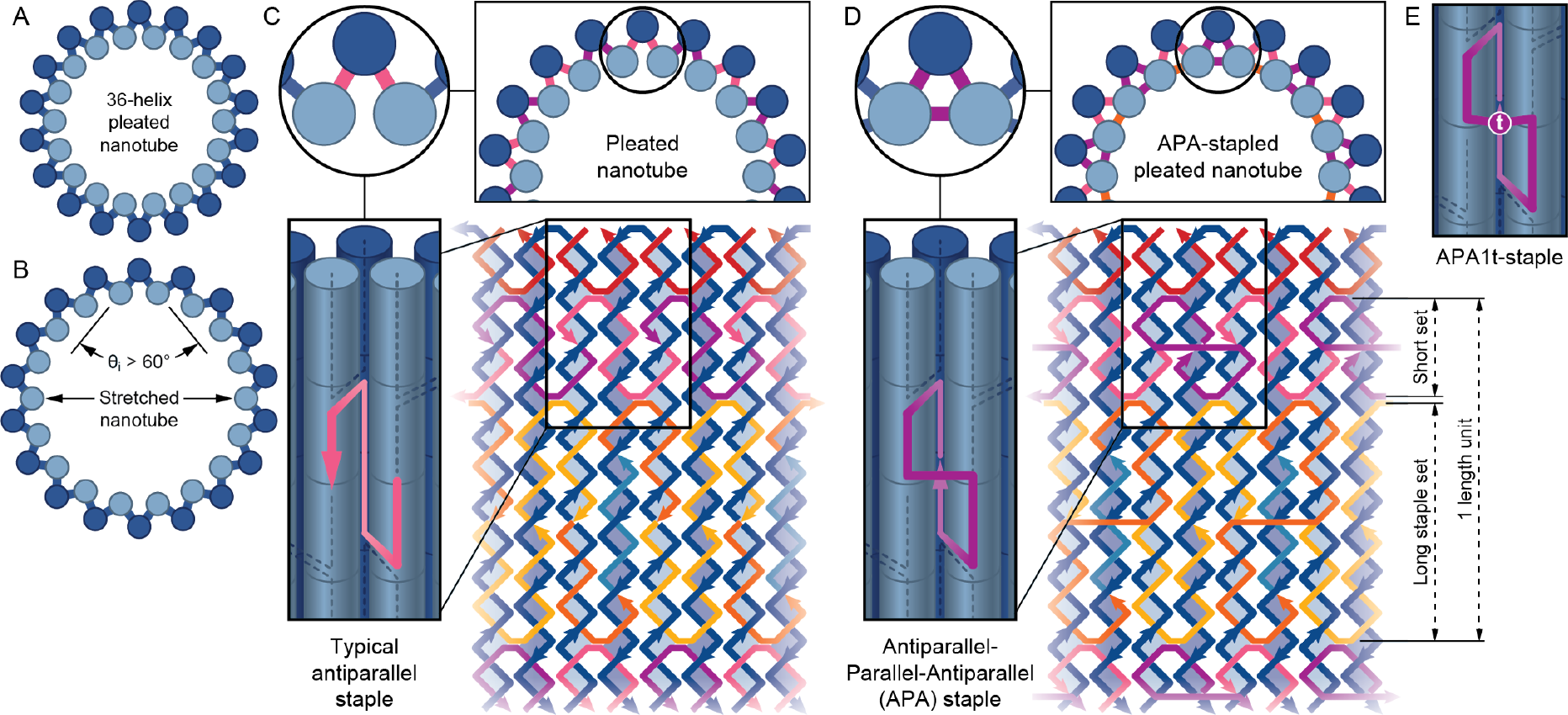
Design and implementation of the APA staple in pleated nanotubes. (A) Schematic of the 36-helix pleated nanotube with internal angle 60°. (B) Schematic of the same nanotube in which electrostatic repulsion between helices on the inner wall result in an internal angle larger than 60°. (C) Top: Detail view of a pleated nanotube schematic as in A with three successive helices enlarged.Crossovers are shown as colored lines that connect the helices. Bottom: Description of the typical “S-shaped” staples that populate the structure, in an isometric pictorial view (left) and in an unrolled staple layout (right). (D) Top: Detail view of an APA-stapled pleated nanotube schematic. The additional parallel crossovers can be seen connecting the inner helices of the nanotube. Bottom: Description of the “figure-8” shaped APA staples that are staggered within the structure, in an isometric pictorial view (left) and in an unrolled staple layout (right). Dashed arrows mark one single length unit, which consists of a long and a short staple set. (E) Pictorial representation of the APA1t staple, which is identical to the APA staple, but with the addition of an unpaired thymine (shown as a “t”) in the parallel crossover.

### Design and implementation of an antiparallel-parallel-antiparallel staple

To further rigidify the structure and constrain the widening of the diameter from electrostatic repulsion between adjacent DNA helices, we implemented a new staple configuration that allows for staple crossovers to occur between all DNA helices in a three-helix bundle. Canonical DNA origami staples in a flat sheet tend to adopt an S-shape as they traverse three sequential helices (4). This is because the template strand runs in opposite directions on adjacent helices, and so staples linking these helices must also change direction at the crossovers (Figure 4C). In contrast, DNA helices in the wall of a pleated nanotube occur in a triangular arrangement where the inner wall of the nanotube is formed by adjacent parallel DNA helices, between which standard antiparallel crossovers do not occur. Nanotube expansion from electrostatic repulsion between inner helices is restrained only by the torsional stiffness of DNA. We therefore introduced a new staple configuration that adopts a “figure 8” shape and forms a single parallel crossover between inner helices in addition to the antiparallel crossovers. Parallel crossovers have previously been shown to be effective in creating DNA origami structures with unidirectional template strand arrangement and in controlling the orientation of DNA origami structures linked by hybridizing staple extensions (64,65). When incorporated into a pleated nanotube, the staple’s path can be described by following its route from the 5’ to 3’ end. The path begins on the outer helix, where it routes via an antiparallel crossover to a neighbouring inner helix, a parallel crossover to a second inner helix and finally via another antiparallel crossover back to the original outer helix (Figure 4D). For this reason, we call the new type of staple the antiparallel-parallel-antiparallel (APA) staple. The APA staple is suitable for use in pleated nanotubes with an internal angle of 60° and for other DNA nanostructures comprised of three-helix bundles. Here we demonstrate its application in pleated DNA origami nanotubes. By linking the helices on the inner surface of the ring, the APA staple should reduce the expansion of the nanotube diameter from electrostatic repulsion resulting in a more rigid structure that more closely matches its design.

APA staples were incorporated into the 36- and 52-helix pleated nanotube designs. Locations for the parallel crossovers on adjacent inner helices were chosen by selecting pairs of C3’ carbon atoms whose distance most closely matched the ideal distance. Examination of these parallel crossover locations revealed three important design considerations for the incorporation of APA staples into pleated nanotubes. Firstly, in order to provide enough base-pairs (> 7) on either side of the parallel crossover for adequate stability, the distance between crossovers had to be increased by a full turn in half of the staples. This reduces the overall number of crossovers and results in a shorter set of staples running in one direction around the nanotube and a longer set running in the opposite direction (Figure 4D). This extra turn had been included in the design of the non-APA-stapled 36- and 52-helix nanotubes so that the effect of the APA staple could be assessed independently of crossover density. The second design consideration was that the routing of APA staples makes them susceptible to form kinetically-trapped configurations that prevent their proper incorporation into the origami structure. This is especially true for the long APA staples, in which tight binding (more than 10 bp) of the 3’ and 5’ ends of the staple to the outer helix may prevent the rest of the staple from weaving around the inner helices of the origami structure (Supplementary Figure S8). To prevent this in the pleated nanotubes, long APA staples were shortened at their 3’ and 5’ ends to reduce their affinity to the β-helix thereby reducing the likelihood of forming a kinetic trap. An additional short DNA strand was used to bind to the resulting unpaired bases in the template strand. Third, for the crossovers to maintain *n/2* symmetry, the staple routing must be identical in every second helix. However, by also satisfying the constraint for crossovers to occur at optimal locations, the search algorithm can construct a physically impossible design where adjacent APA staples overlap or bind to the same template bases (Supplementary Figure S9). As a practical solution to this, parallel crossovers were staggered between every second pair of helices resulting in staple routing with *n/4* rotational symmetry.

We synthesized the 36- and 52-helix pleated nanotubes with APA staples by replacing half of the short regular staples and half of the long regular staples with APA staples as illustrated in Figure 4. In measurements taken from cryo-EM images, widths of pleated nanotubes with APA staples were less than nanotubes without APA staples (Figure 5A; Supplementary Figures S6 and S7), indicating that the APA staple reduces the expansion of nanotubes from the electrostatic repulsion between helices. This was also evident in nanotube width measurements in both TEM (Figure 5B) and AFM (Supplementary Figure S5). The APA staple also appeared to make the 36-helix pleated nanotube more rigid and resilient to flattening. This was evident in the both the TEM and AFM micrographs. In the TEM images, the width of APA-stapled nanotubes appeared not only narrower than the non-APA-stapled nanotubes (Figure 5B), but had more prominent walls and more easily accumulated stain in the centre of the nanotube (Figure 5C). Both are indicative of a more raised and hollow structure. In addition, AFM micrographs show that well-formed 36-helix nanotubes with APA staples had a more uniform and greater height compared with nanotubes without APA staples (Supplementary Figure S5A and B). This demonstrates that APA-stapled nanotubes were also more resilient to flattening from the AFM tip. However, these nanotubes were also more susceptible to breaking, with 37.6% of APA-stapled 36-helix nanotubes appearing as unrolled structures when imaged with AFM. Consistent with this are images of APA-stapled 52-helix pleated nanotubes, where 73% of nanotubes appeared unrolled when imaged with AFM and 56% appeared to have split in TEM micrographs (Figure 5B). End-on images of these nanotubes that adsorbed to the carbon support of the cryo-EM grid illustrate how these split nanotubes form a ‘scroll-like’ structure with higher curvature (Supplementary Figure S7E). Thus, while the APA staples do rigidify pleated nanotubes and allow for better control of their diameter, APA staples also appear to contribute additional strain on the nanotubes that makes them more susceptible to splitting.

**Figure 5.**
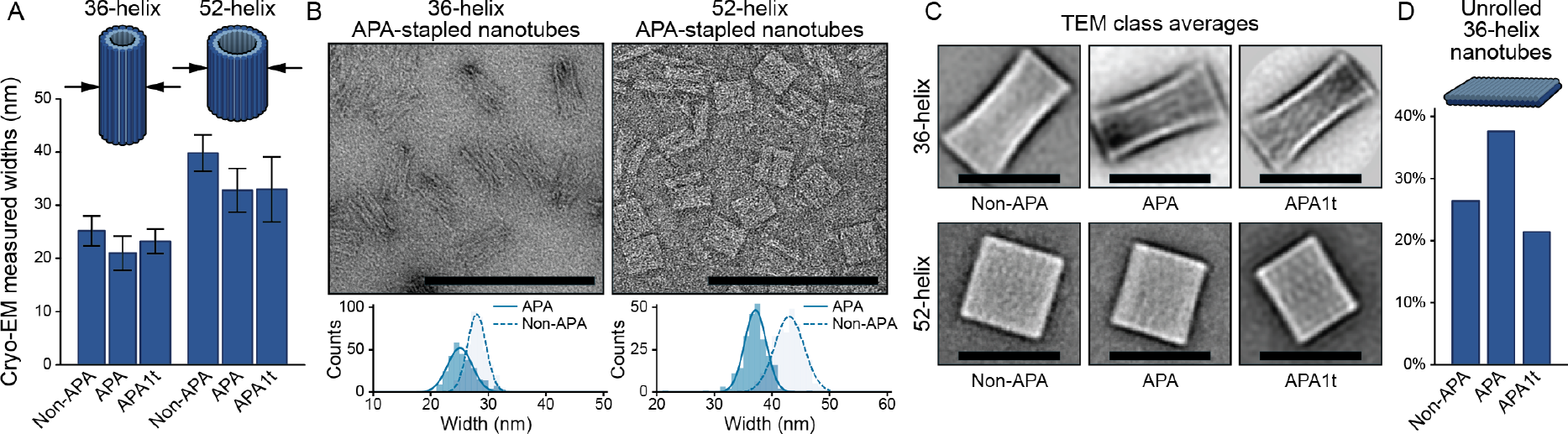
Dimensions, TEM imaging and stability of APA-stapled DNA origami nanotubes. (A) Bar chart of nanotube widths measured from cryo-EM micrographs. Error bars represent standard deviation. (B) Top: Representative TEM micrographs of 36-helix (left) and 52-helix (right) APA-stapled pleated nanotubes. Scale bars in micrographs represent 200 nm; Bottom: Histograms of measurements comparing the widths of APA-stapled (solid) and non-APA stapled (dashed) pleated nanotubes. (C) Comparison of class averages of nanotubes in TEM micrographs. In each case the class average shown is the one with the most particles assigned to that class. Scale bars represent 50 nm. (D) Comparison of percentage of 36-helix nanotubes that were ‘unrolled’ in the AFM micrographs.

One reason for the additional strain may be because the distance between C3’ carbon atoms at parallel crossovers was substantially larger than the ideal distance. This would result in strain at these crossovers and hence potential bending, twisting or shearing of the helices, or even the breaking of hydrogen bonds at adjacent base-pairings. Therefore, to alleviate this additional strain and make APA-stapled nanotubes less susceptible to splitting, we incorporated a single additional thymine base at the parallel crossover location in all APA staples (Figure 4E; Supplementary Figure S10). The incorporation of this “APA1t” staple lowered the proportion of unrolled 36-helix nanotubes visualized with AFM (21.4%) to be similar to the proportions of unrolled structures seen in nanotubes without APA staples (Figure 5D). Moreover, there were substantially fewer split 52-helix nanotubes (33%) with APA1t staples compared to those with APA staples, when seen in TEM images. Additionally, for both the 36- and 52-helix nanotubes, the incorporation of the APA1t staple did not significantly change the measured cross-sectional widths when compared with APA-stapled nanotubes. This was seen both in solution (Figure 5A; Supplementary Figure S8) and on the TEM grid surface (Figure 5C; Supplementary Table S1). These data confirm that the addition of a thymine base at the parallel crossover in APA staples can reduce strain-induced breakages without affecting the average geometry of the structure. Moreover, we suggest that the incorporation of additional thymine bases at the parallel crossover could provide a means to finely tune the internal angle and hence the diameter of pleated DNA origami nanotubes.

Here we report a simple algorithm that comprehensively searches the design space and provides the ideal crossover locations for the construction of minimally-strained DNA origami nanotubes. This provides a facile approach for the construction of nanotubes with an arbitrary even number of helices as well as the construction of nanotubes with a pleated wall. Pleated nanotubes reduce the flexibility inherent in single-layered origami structures, allow for a continuous range of nanotube diameters and interhelical distances between parallel helices, which we demonstrate can be arranged side-by-side. Characterization of pleated nanotubes revealed that these were subject to expansion from electrostatic repulsion between adjacent parallel helices because typical DNA origami staples do not crossover between parallel helices. We therefore also introduce a new staple routing, which we have named the APA staple, which includes a crossover between parallel helices. The APA staple prevents nanotube expansion allowing the construction of pleated nanotubes that closely fit their design. Moreover, by incorporating additional thymine bases at the crossover location, APA staples provide a simple mechanism to tune the diameter of the nanotube by replacing only a small subset of DNA staples. Combined these findings provide an accessible approach for the design and synthesis of DNA origami nanotubes and increase the design space to allow for a continuous range of interhelical angles, diameters and interhelical spacing between arbitrary numbers of radially symmetric parallel helices.

## ACKNOWLEDGEMENT

The authors thank the facilities of Microscopy Australia at the Electron Microscope Unit, Mark Wainwright Analytical Centre, UNSW Sydney. We would also like to thank Professor Keiichi Namba for providing access to cryo-EM facilities at the Graduate School of Frontier Biosciences, Osaka University, Japan, as well as Dr Rhiannon Kuchel at UNSW Electron Microscope Unit for preliminary cryo-EM imaging, and Dr Julian C. Berengut at the UNSW School of Physics for assistance in writing the algorithm.

