## Supplementary material for "Design and Synthesis of Pleated DNA Origami Nanotubes with Adjustable Diameters": Figure S

---

**Supplementary Note 1 - Search algorithm for locating minimally-strained crossovers.**

*Determination of DNA backbone coordinates.* The backbone of a DNA double helix was modelled geometrically with a parametric equation for helices:

$$\left\{r\sin\left[t+\left(\frac{\pm g}{2}\right)\circ\right], -r\cos\left[t+\left(\frac{\pm g}{2}\right)\circ\right], \frac{t\times pitch_{nm}}{\pi}\right\}$$

where $r$ is the radius of an ideal DNA helix (1.0 nm), $g$ is the minor groove angle of the DNA (135°) and is positive for one strand of DNA and negative for the complementary strand, and $pitch_{nm}$ is the helical pitch of a DNA helix (3.55 nm). The coordinates of C3' carbon atoms along the helical path were defined by the helical pitch of DNA on a double helix in base pair units ($pitch_{bp} = 10.5\ bp/turn$) at points where $t = base_i \times \left(\frac{2\pi}{pitch_{bp}}\right)\circ$ where $i$ is the base number and ranges from 0 to the length of the DNA helix in base pair units.

Thus, a set of coordinates for C3' carbon atoms and NEMids could be determined for a DNA helix. This set of coordinates was transformed to construct an arrangement of three helices with the desired internal angle (helix α, helix β and helix α+1), (Figure 1D and 1E) and an inter-helix distance of 2.25 nm. This corresponds to a geometry where perfectly-aligned crossovers should have an ideal bond-distance. For a given internal angle and number of helices in the nanotube, the minimally strained crossover locations were determined for all combinations of $\theta_\alpha$ and $\theta_\beta$ angles (Figure 1D and E), which were sampled every 1°.

*Identification of crossover locations.* From these geometric models, all inter-NEMid and inter-C3' carbon distances and strain scores can be determined. To reduce the number of calculations and ensure an average staple crossover density of one crossover every 1.5 turns of a DNA helix, it was useful to first approximate the location of staple crossovers. This was done as follows: Template crossovers are placed at ends of DNA helices in the nanotube. At the 'bottom' end of the nanotube, the algorithm first determines whether the template crosses over between helix α and helix β, or between helix β and helix α+1 by assessing the strain score of each. The location of the template crossover at the bottom of helix β then sets a reference point from which the location of staple crossovers that form the first length unit can be approximated. If the first template crossover occurs between helix β and helix α, then the first staple crossover will occur between helix β and helix α+1 ($\beta \rightarrow \alpha + 1$) around where:

$$t_{\beta\rightarrow\alpha+1} = 2\pi - g - \theta_i$$

and the second staple crossover between helix β and helix α ($\beta \rightarrow \alpha$) occurs at:

$$t_{\beta\rightarrow\alpha} = t_{\beta\rightarrow\alpha+1} + \theta_i + 2\pi$$

Alternatively, if the first template crossover occurs between helix β and helix α+1, then the first staple crossover will occur between helix α and helix β around where:

$$t_{\beta\rightarrow\alpha} = 2\pi - g + \theta_i$$

and the second staple crossover between helix β and helix α+1 occurs at:

$$t_{\beta+1\rightarrow\alpha} = t_{\beta\rightarrow\alpha} - \theta_i + 2\pi$$

The remaining crossovers between helix β and each of its neighboring DNA helices occur every three turns and are staggered between staples that crossover to helix α and helix α+1. This pattern is repeated for the number of desired length units. Exceptions to this staple patterning occurred either when the distance between the template and the nearest staple crossover was less than seven base pairs, in which case the staple was extended by a full turn, or to incorporate APA staples in pleated nanotubes where every second staple was also extended by one full turn. These approximate crossover locations are then used to create a search space within four base-pairs of the approximate locations. All NEMid pairs within each search space are then measured and converted to a strain score, and the pair with the lowest score is used to determine the minimally-strained crossover at that location. The staple crossover at the top end of nanotube is approximated in a similar way after the last staple crossover. Overall strain scores were defined as the sum of the squared difference between measured and ideal inter-NEMid and inter-template crossover distances. Subsequently, the crossover locations with the minimal strain score can result in a difference in height of 1-2 base pairs in every second DNA helix. The search algorithm was implemented in Wolfram Mathematica version 11.2. Once minimally-strained crossover locations were identified, these were entered manually into Cadnano v2 (1) to determine staple sequences.

*Details of nanotube designs that were synthesized.* In some cases, the crossover scheme chosen for the synthesized nanotube design was not the lowest scoring solution as output by the algorithm. For example, the 20-helix design selected for synthesis has a similarly low score as the number one solution, but a slightly different template crossover at one end of the nanotube which was more conducive to subsequent experiments. The algorithm outputs and an explanation of the chosen design are shown in Figure S12. Crossover designs were entered manually into Cadnano (schematics shown in Figure S13-S18) to calculate DNA staple sequences, which are available in an Excel spreadsheet in the supplementary materials.

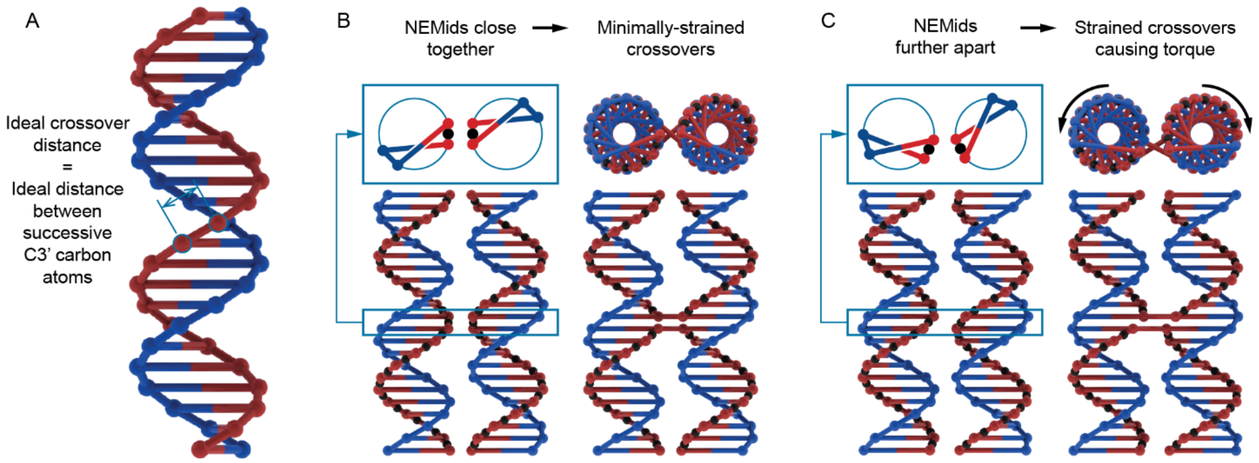


**Figure S1. Calculation of staple crossover strain.** (A) A depiction of a DNA double-helix, with the locations of C3' carbon atoms shown as red or blue spheres. The ideal distance between C3' carbon atoms on either side a crossover span was assumed to be the same as the distance between successive C3' carbon atoms in an idealized double helix. This was calculated to be approximately 0.68 nm. (B) Two adjacent double helices viewed from the front. The staple strand C3' carbon atoms are shown as red spheres. The NEMids are the points midway between them and are shown as black spheres. When two NEMids are the ideal distance apart, the crossovers formed are also ideal, resulting in minimal strain to the helices. (C) If the inter-NEMid distance is not ideal, the resulting crossovers above and below the NEMid will be strained. This was quantified as the sum of the squared difference between measured and ideal inter-NEMid distances.

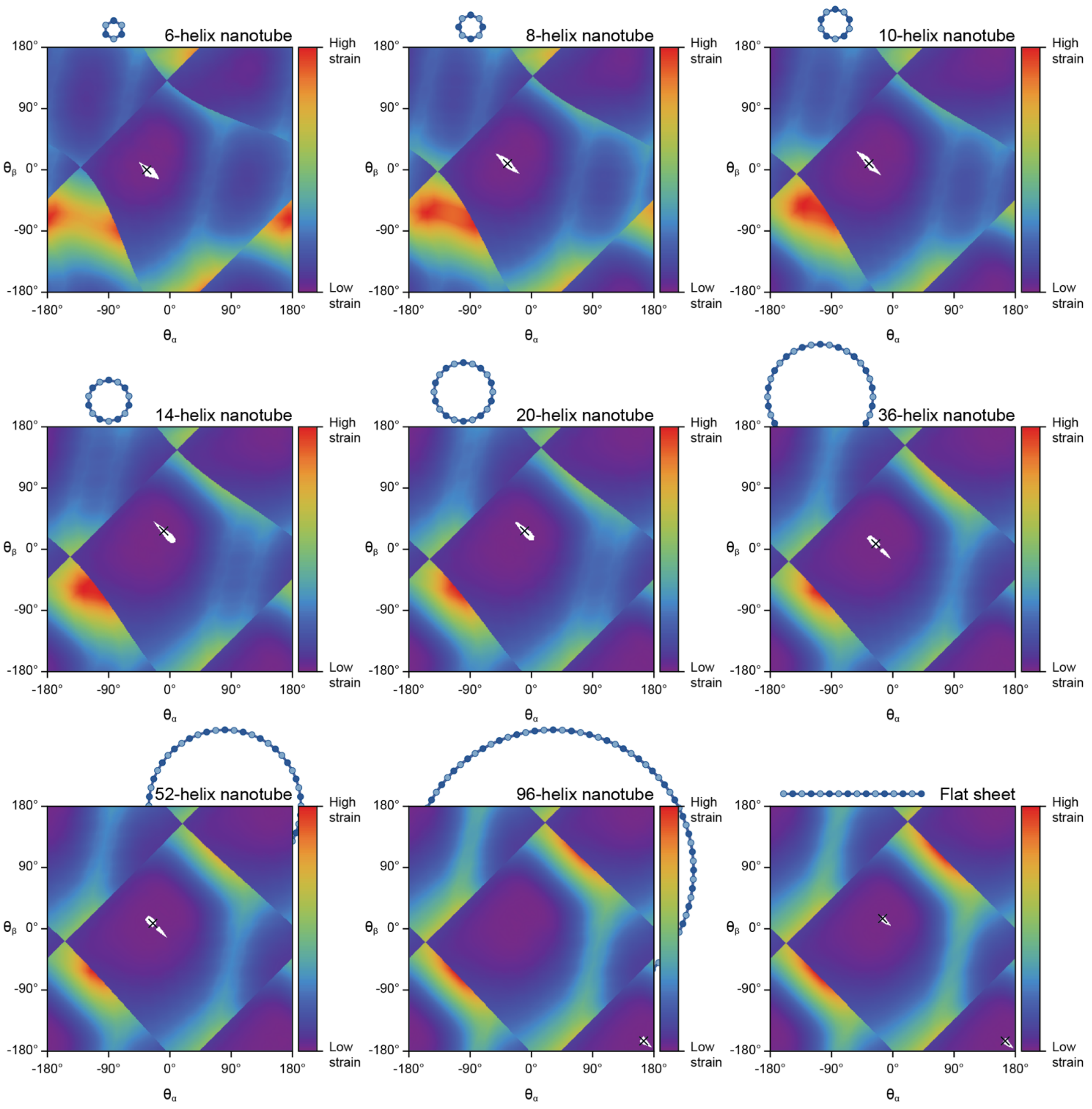


**Figure S2. Graphs of strain scores calculated for different nanotube sizes.** $\theta_\alpha$ refers to the rotation angle of helices α and α+1, and $\theta_\beta$ refers to the rotation angle of helix β. Scores correspond to the set of crossover locations where crossover geometry was closest to ideal for each combination of rotational angles. Maximum and minimum scores were normalized for comparison and colored as a spectrum. All length units were set to 6. In each design, the helix orientations that resulted in the lowest scoring solution is marked with a black "X". Often, the same set of crossover locations represented the minimal strain solution to several helical orientations. Thus, all helical orientations for which the lowest scoring solution was found are shown in white. In the case of the flat sheet design, two opposite solutions had equally low strain scores.

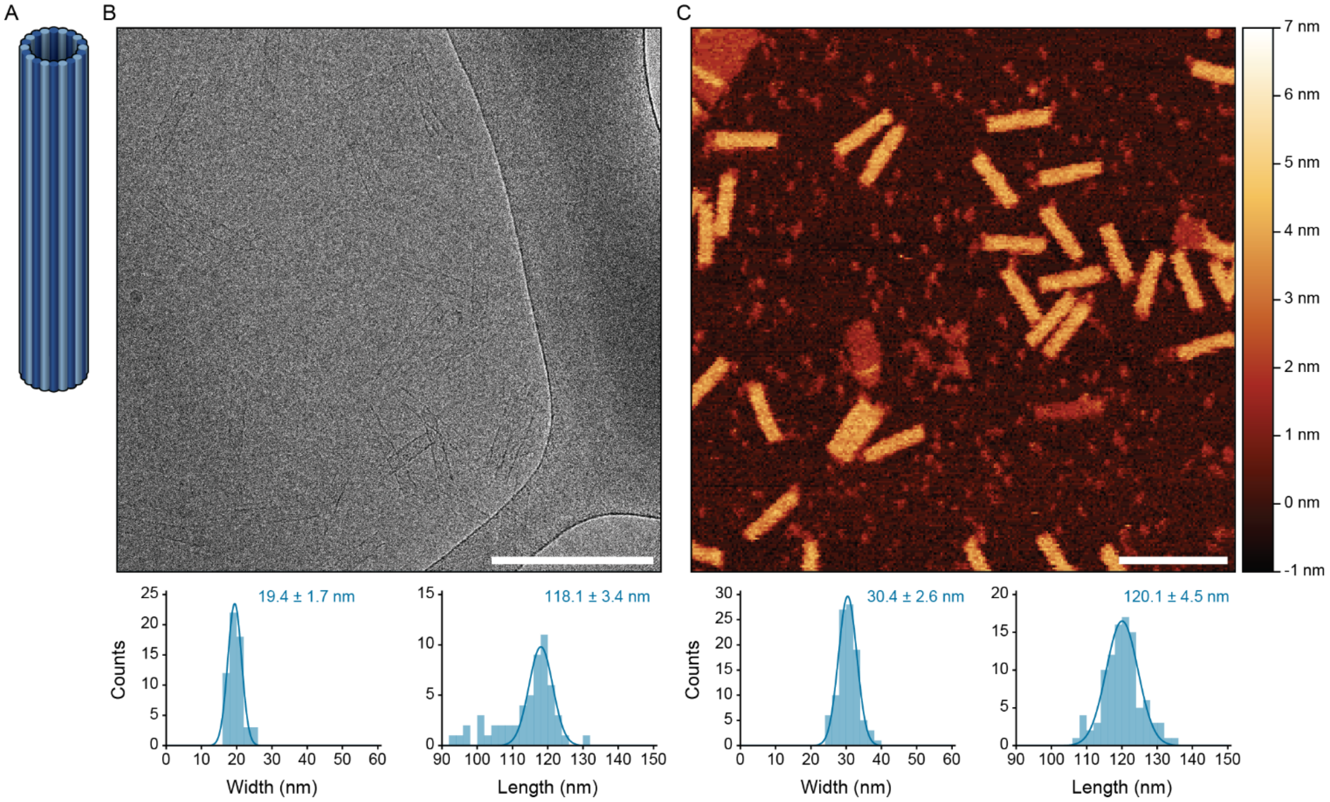


**Figure S3. Cryo-EM and AFM imaging of the regular 20-helix nanotube.** (A) Rendering of the 20-helix nanotube. (B) Top: Representative cryo-EM micrograph (Scale bar is 200 nm) and Bottom: Histograms of measured of widths and lengths or nanotubes imaged with cryo-EM. (C) Top: Representative AFM micrograph (Scale bar is 200 nm); Bottom: Histograms of measured of widths and lengths of nanotubes imaged with AFM.

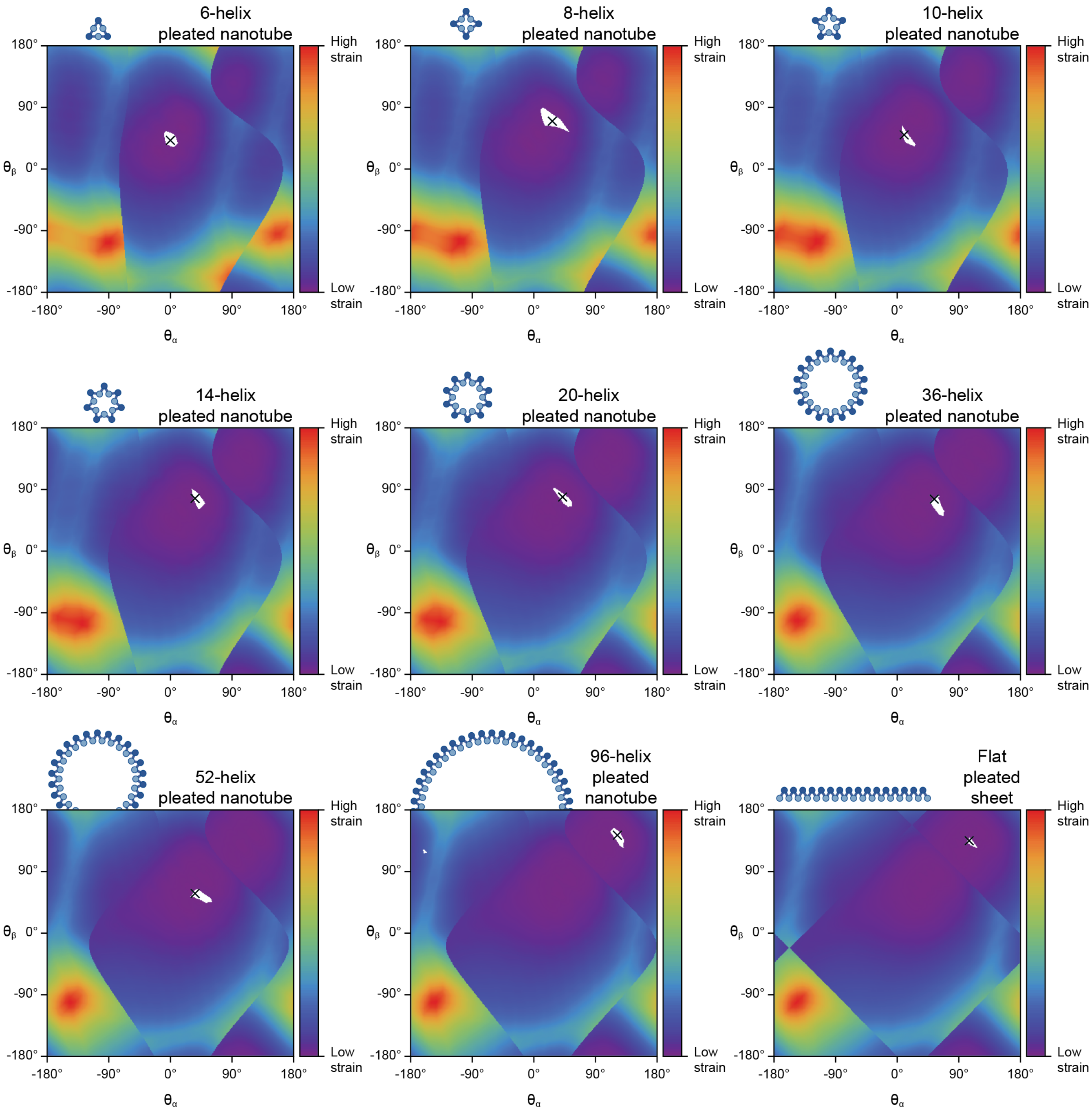


**Figure S4. Graphs of strain scores calculated for different pleated nanotube sizes.** $\theta_\alpha$ refers to the rotation angle of helices α and α+1, while $\theta_\beta$ refers to the rotation angle of helix β. Scores correspond to the set of crossover locations where crossover geometry was closest to ideal for each combination of rotational angles. Maximum and minimum scores were normalized for comparison and colored as a spectrum. All length units were set to 6. In each design, the helix orientations that resulted in the lowest scoring solution is marked with a black "X". All helical orientations for which the lowest scoring set of crossover locations was found are shown in white.

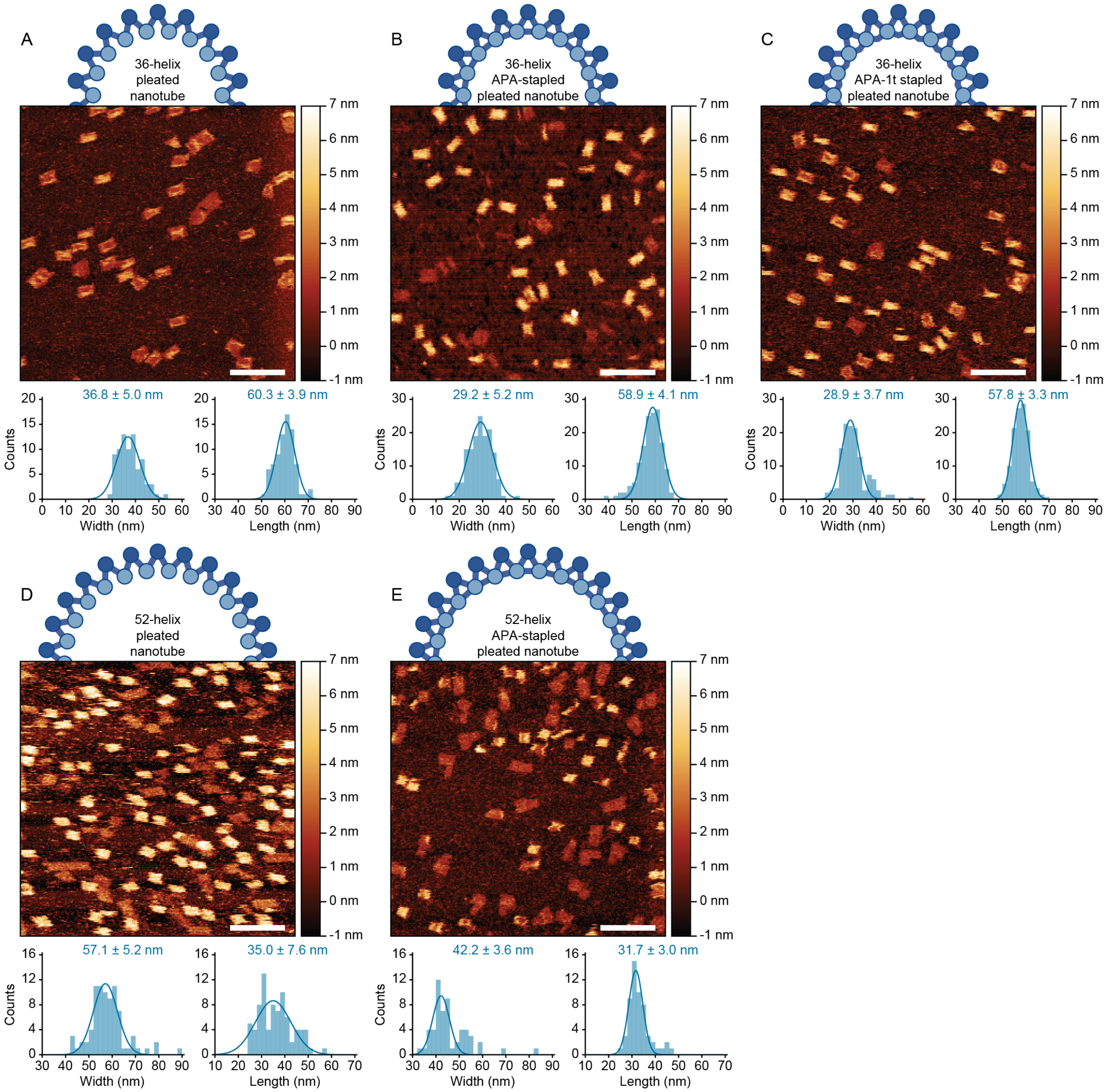


**Figure S5. AFM micrographs of pleated nanotubes.** Schematic of nanotube (top), representative micrograph (middle) and histograms of width and length measurements (bottom) of 36-helix pleated nanotubes (A), 36-helix pleated nanotubes with APA staples (B), 36-helix pleated nanotubes with APA1t staples (C), 52-helix pleated nanotube (D), 52-helix pleated nanotube with APA staples (E).

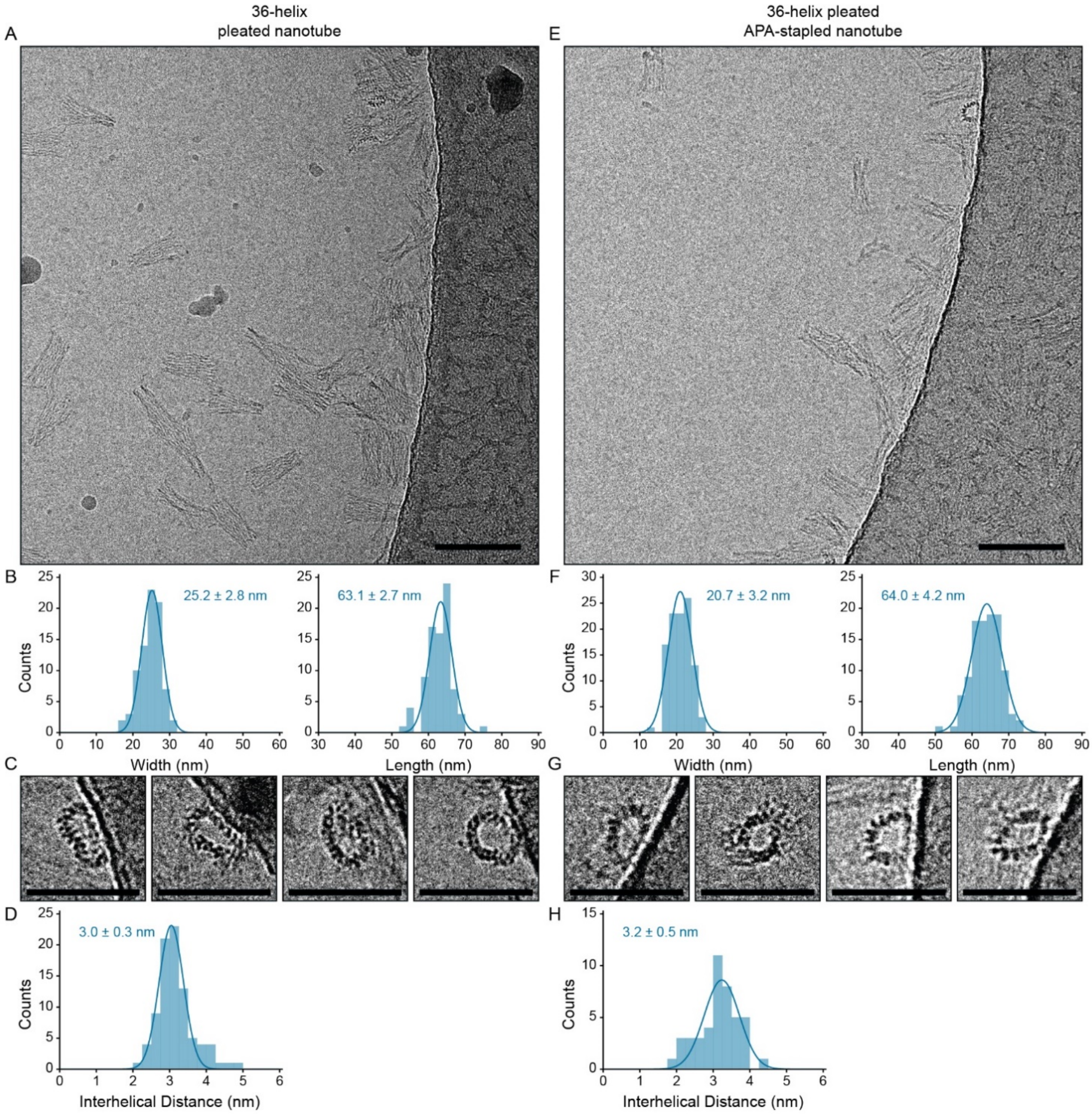


**Figure S6. Cryo-EM micrographs and analysis of 36-helix nanotubes.** (A) Representative cryo-EM micrograph of 36-helix pleated nanotubes (scale bar: 100 nm). (B) Histograms of width and length measurements. (C) Selected end-on views of intact 36-helix pleated nanotubes (scale bars: 50 nm). (D) Histogram of interhelical distance measurements. (E) Representative cryo-EM micrograph of 36-helix APA-stapled pleated nanotubes (scale bar: 100 nm). (F) Histograms of width and length measurements. (G) Selected end-on views of intact 36-helix APA-stapled pleated nanotubes (scale bars: 50 nm). (H) Histogram of interhelical distance measurements.

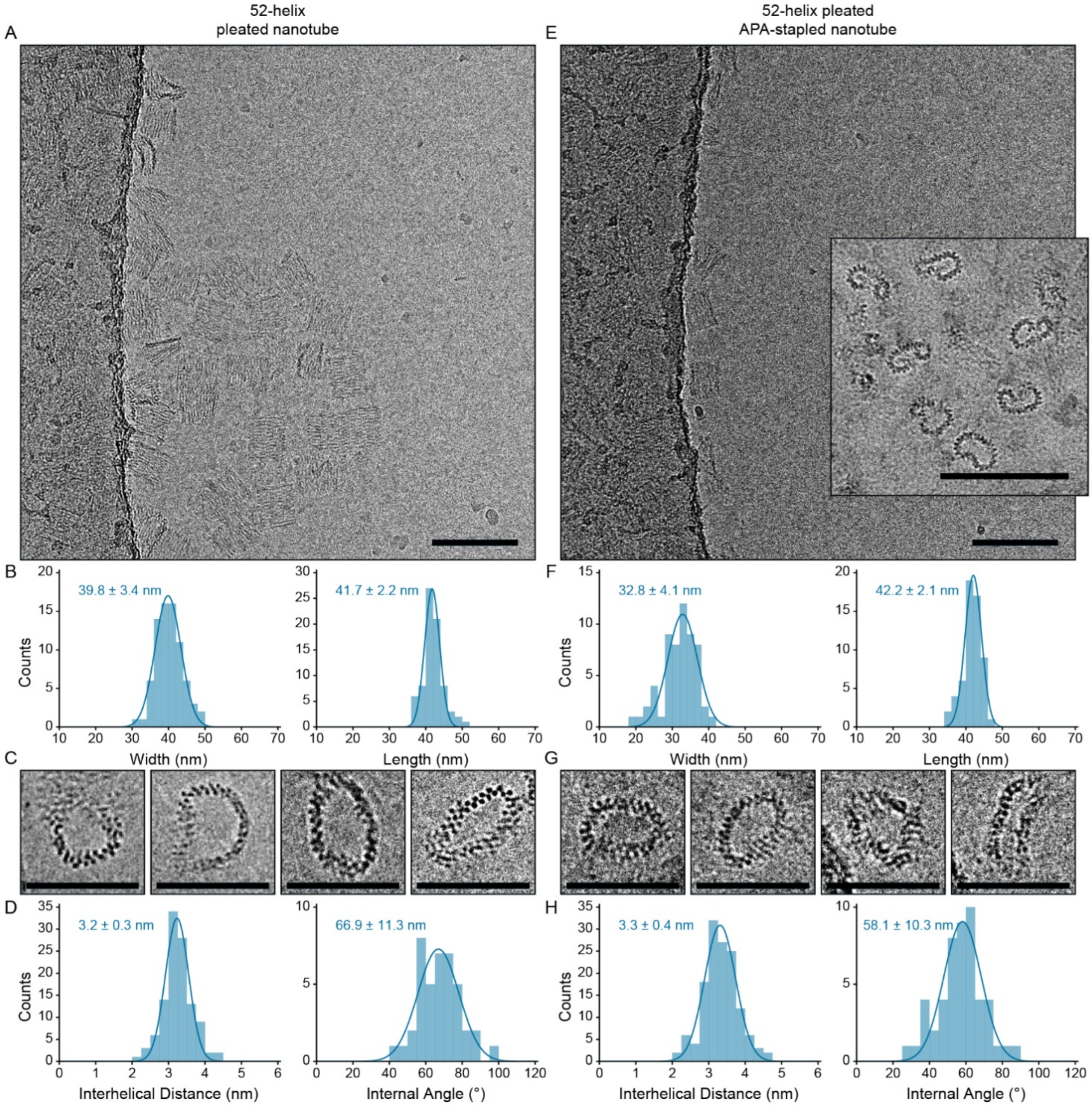


**Figure S7. Cryo-EM micrographs and analysis of 52-helix nanotubes.** (A) Representative cryo-EM micrograph of 52-helix pleated nanotubes (scale bar: 100 nm). (B) Histograms of width and length measurements. (C) Selected end-on views of intact 52-helix pleated nanotubes (scale bars: 50 nm). (D) Histogram of interhelical distance and internal angle measurements. (E) Representative cryo-EM micrograph of 52-helix APA-stapled pleated nanotubes (scale bar: 100 nm). Inset: End-on views of curled APA-stapled pleated nanotubes attached to the carbon support (scale bar: 100 nm). (F) Histograms of width and length measurements. (G) Selected end-on views of intact 52-helix APA-stapled pleated nanotubes (scale bars: 50 nm). (H) Histogram of interhelical distance and internal angle measurements.

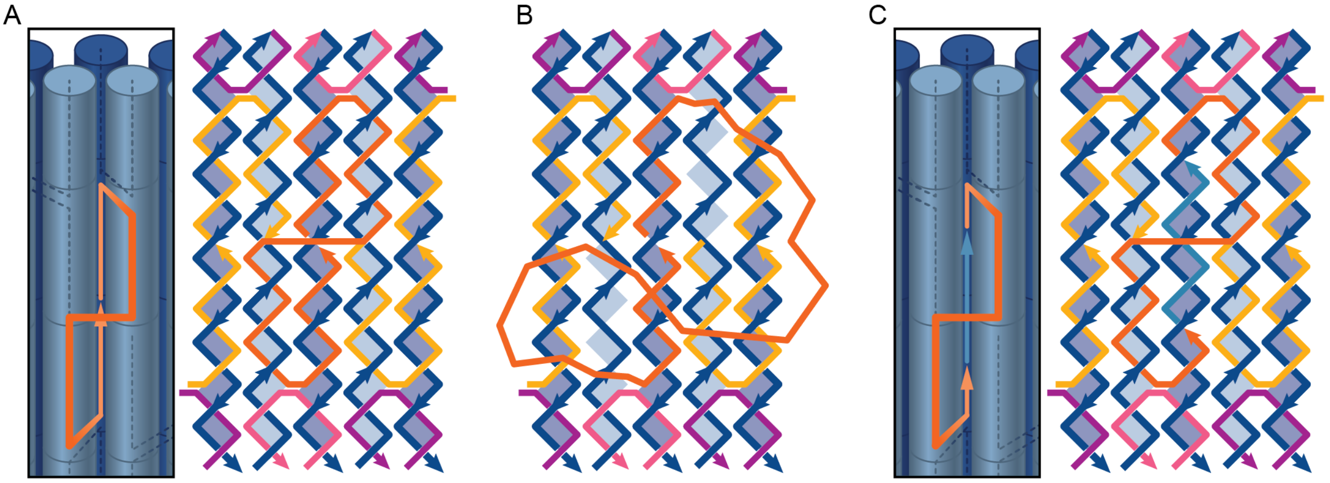


**Figure S8. Avoiding kinetic traps in the long APA staple.** (A) Pictorial view and unrolled staple layout depicting successive helices in an APA-stapled pleated nanotube. The outer helices are depicted as dark blue and the inner helices as light blue. The strand routing of the long APA staple is shown as an orange line with an arrow at the 3' end. Correct incorporation of this staple relies on the 5' and 3' ends of the staple binding to adjacent sections of the scaffold strand of the outer helix, while the rest of the staple weaves around the scaffold that makes up the two inner helices. (B) If the sections that are bound to the outer helix are bound too tightly, we propose that kinetic traps may prevent the middle of the staple from correctly binding. (C) To prevent this, we shorten both ends of the APA staple to reduce affinity to the outer helix and facilitate strand migration. The gap created is then filled with a short filler strand, shown in light blue.

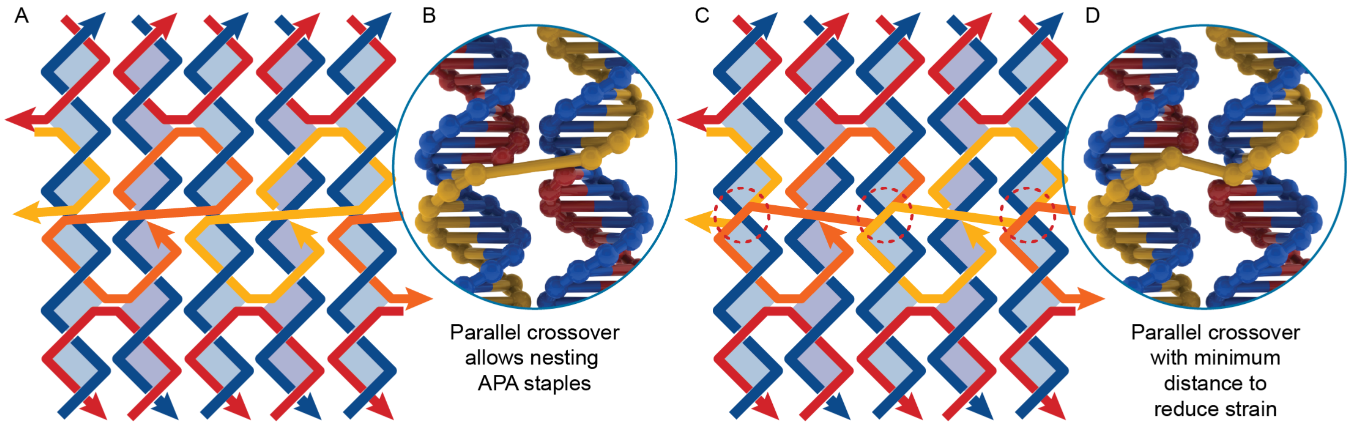


**Figure S9. How the choice of parallel crossover position affects strain and radial symmetry.** (A) In order to gain full *n*/2 rotational symmetry, the APA staples (yellow and orange) must 'nest' side-by-side so that every second helix is identical. (B) In many cases, this is only possible by including a very long parallel crossover, which would likely induce strain into the structure. (C,D) If the length of the parallel crossovers is optimized, overlaps occur (circled in red dotted lines) where successive APA staples need to bind to the same bases of the template strand. Areas of overlap are circled with a red dotted line.

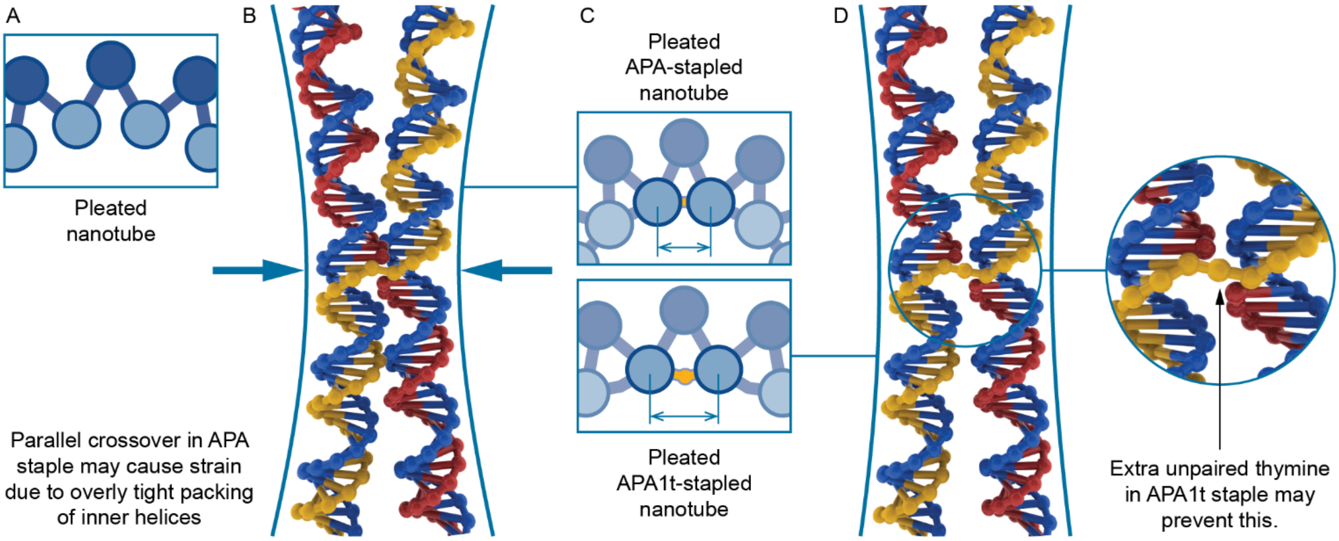


**Figure S10. Description of the APA1t staple.** (A) Typical layout of a section of pleated nanotube, with the outer helices depicted as dark blue circles, inner helices depicted as light blue circles, and crossovers depicted as blue lines connecting the helices. (B) Implementation of the APA staple involves a parallel crossover between inner helices. The distance that this crossover must span is generally greater than the ideal crossover distance, which may result in the helices being pulled together in such a way as to introduce strain into the structure. (C) Layout diagrams of sections of APA-stapled (top) and APA1t-stapled (bottom) nanotubes with arrows indicating the interhelical distance between inner helices. (D) The APA1t staple includes an unpaired thymine in the parallel crossover, which was added to relieve the strain in B.

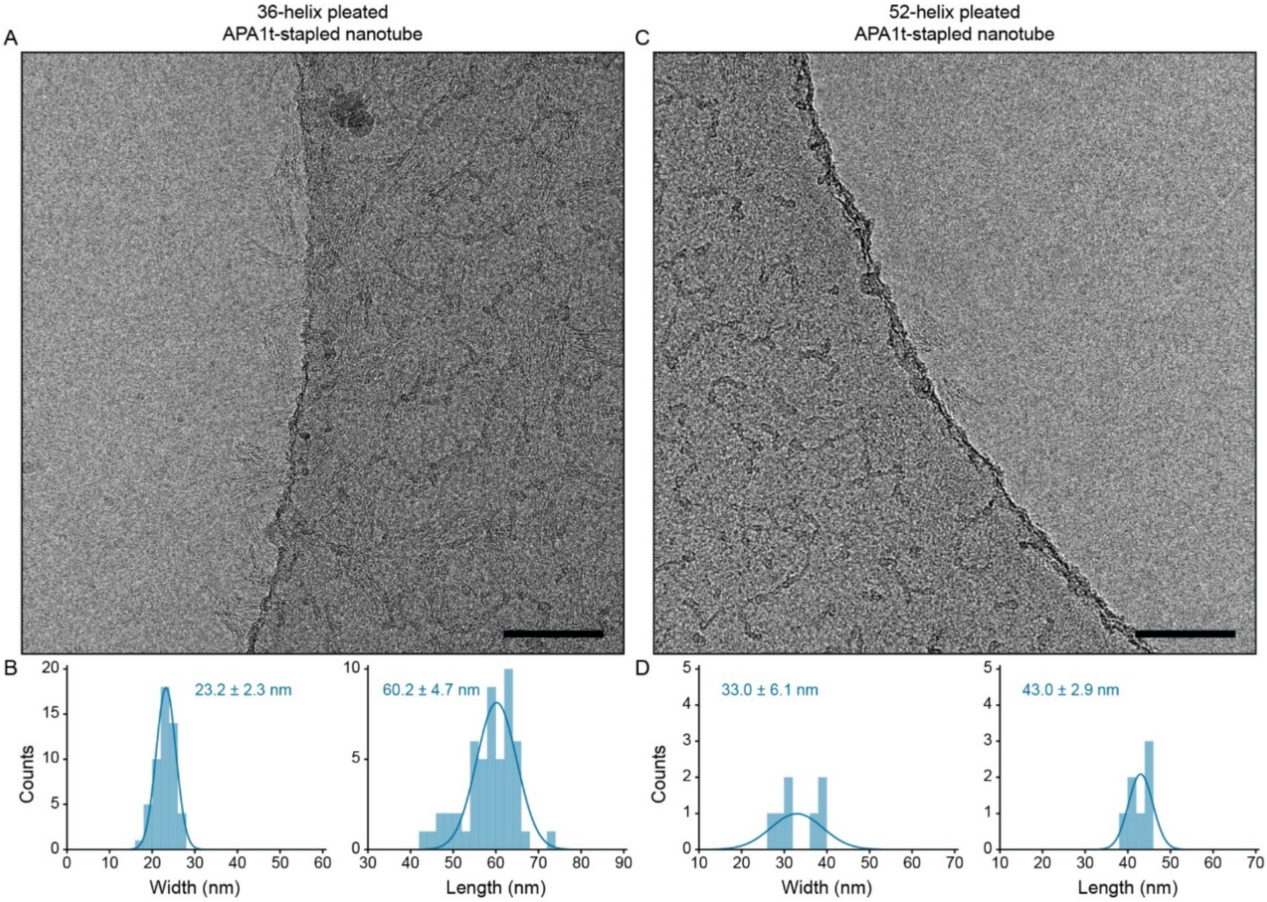


**Figure S11. Cryo-EM micrographs and analysis of APA1t-stapled pleated nanotubes**. (A) Representative cryo-EM micrograph of 36-helix APA1t-stapled pleated nanotubes (scale bar: 100 nm). (B) Histograms of width and length measurements. (C) Representative cryo-EM micrograph of 52-helix APA1t-stapled pleated nanotubes (scale bar: 100 nm). (D) Histograms of width and length measurements.

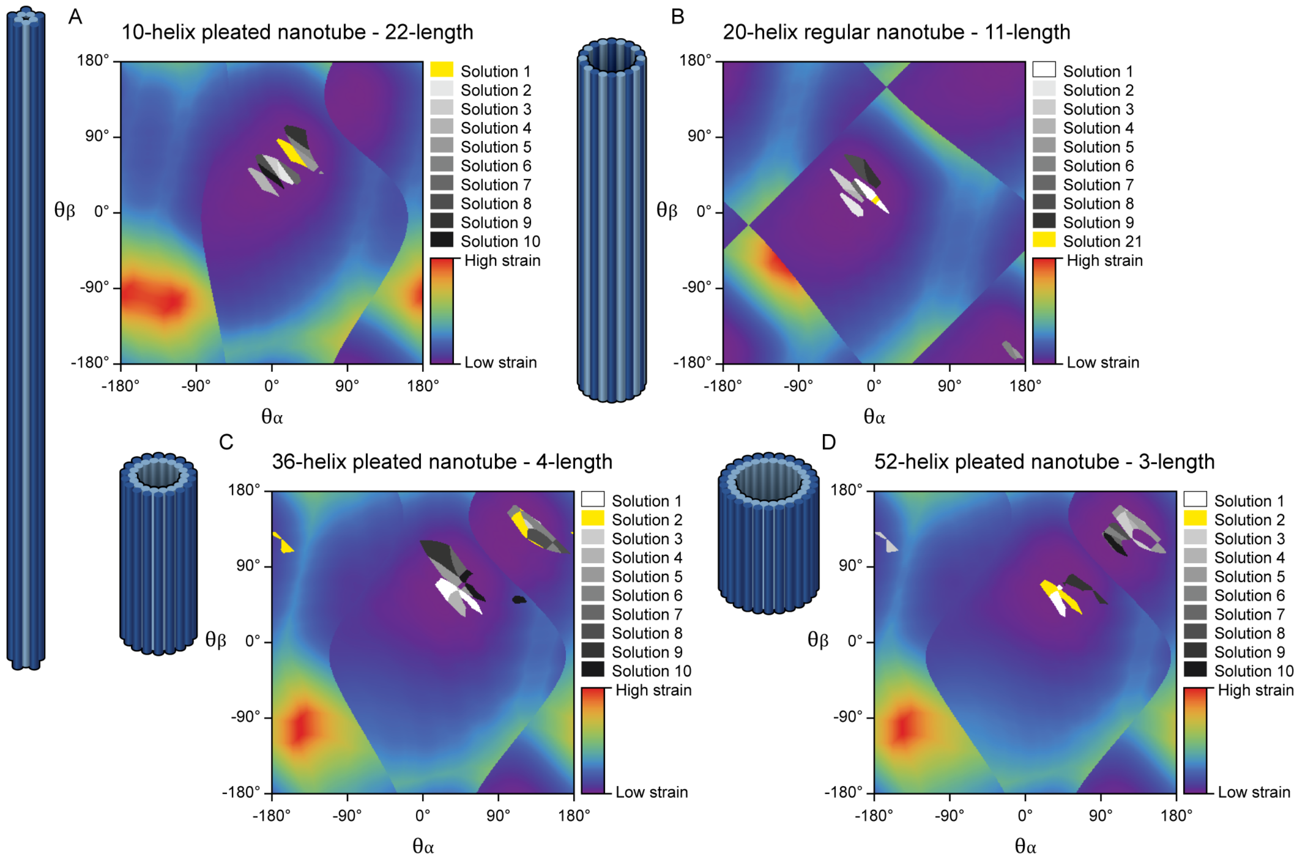


**Figure S12. Output from the algorithm with parameters of nanotubes we synthesized.** Plots of strain scores for the 10-helix pleated nanotube (A), the 20-helix regular nanotube (B), the 36-helix pleated nanotube (C) and the 52-helix pleated nanotube (D). The best ten crossover solutions are plotted in shades of grey in all the angles in which they occurred and the design selected for synthesis and characterization is in yellow. For the 20-helix and 52-helix nanotubes, the solution that was synthesized and characterized had the same staple crossover positions as the lowest scoring solution and similarly low score, but slightly different scaffold crossover positions at one end of the nanotube. For the 36-helix pleated nanotube, the strain score of the synthesized solution was also similar to the lowest scoring solution but the scaffold strand was running in the opposite direction.

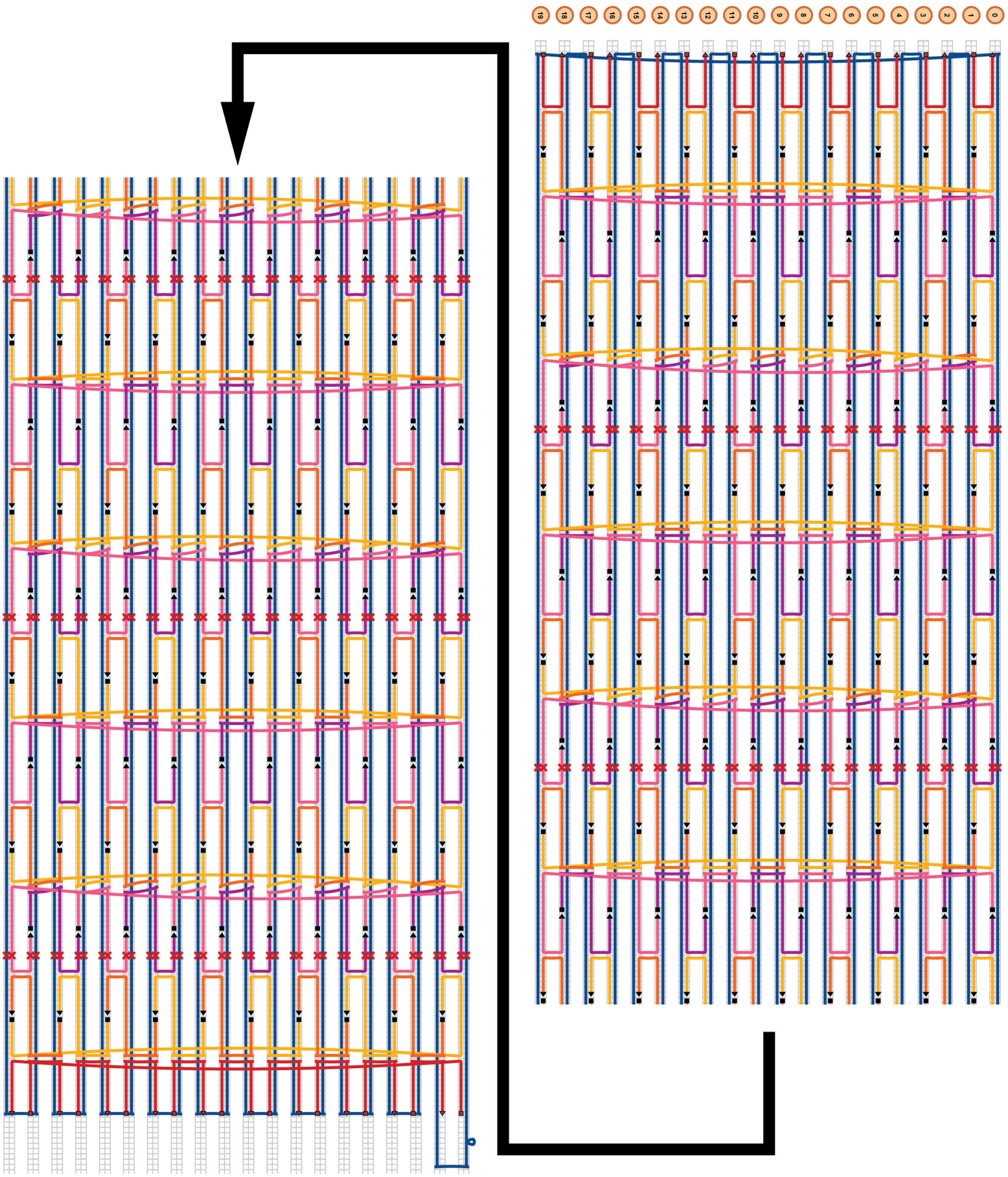


**Figure S13. Cadnano schematic of the 20-helix nanotube.**

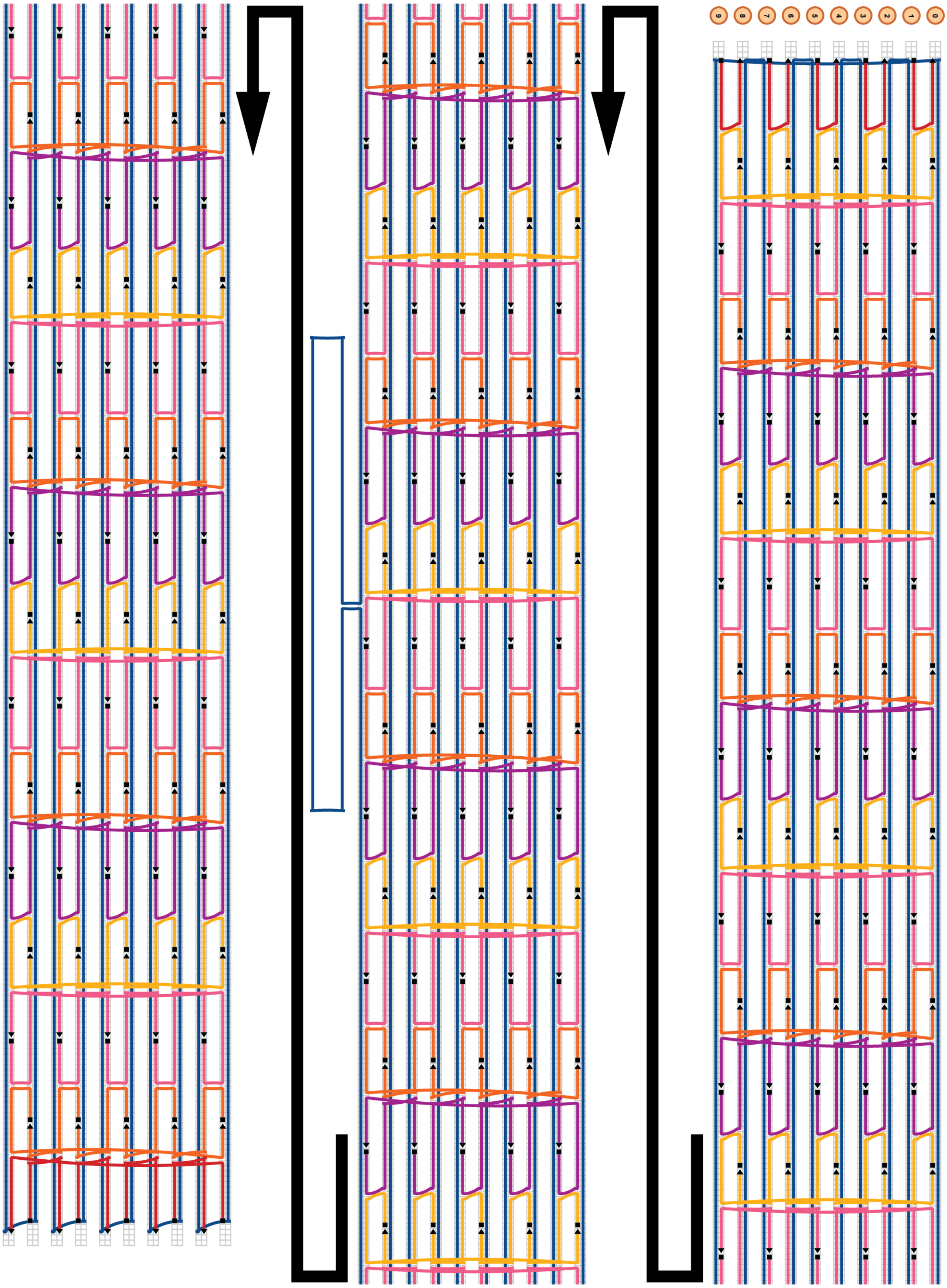


**Figure S14. Cadnano schematic of the 10-helix pleated nanotube.**

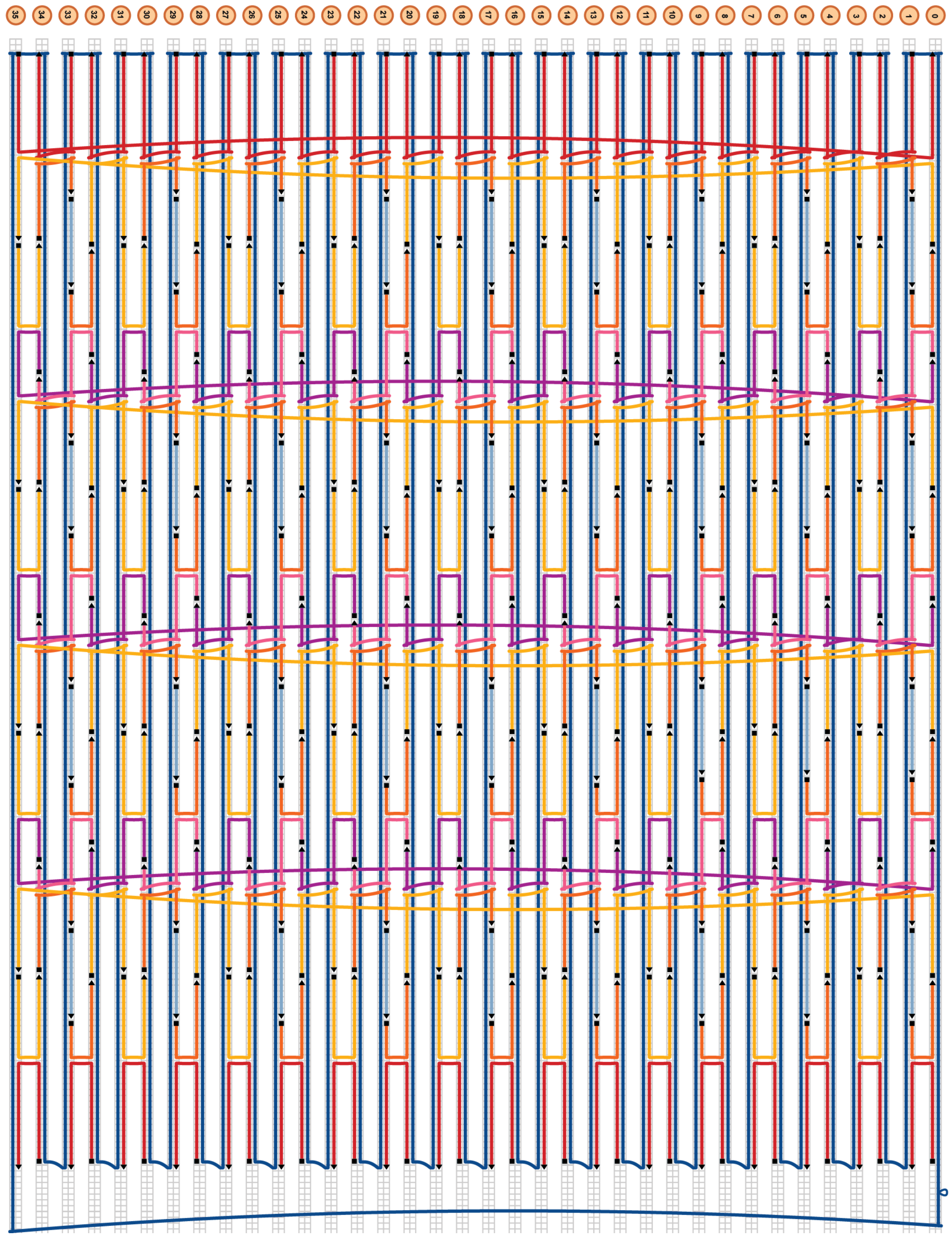


**Figure S15. Cadnano schematic of the 36-helix pleated nanotube.**

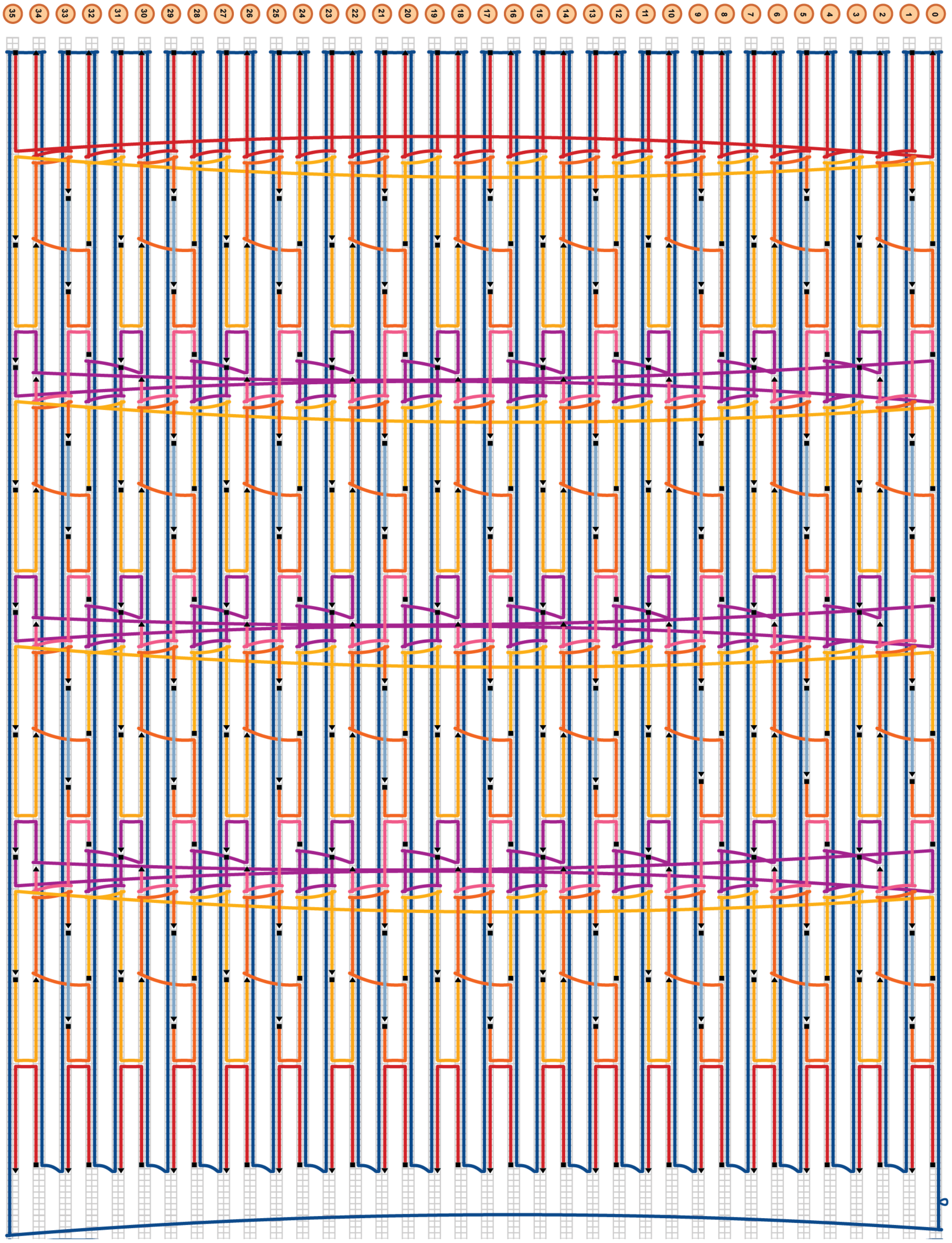


**Figure S16. Cadnano schematic of the 36-helix pleated APA-stapled nanotube.**

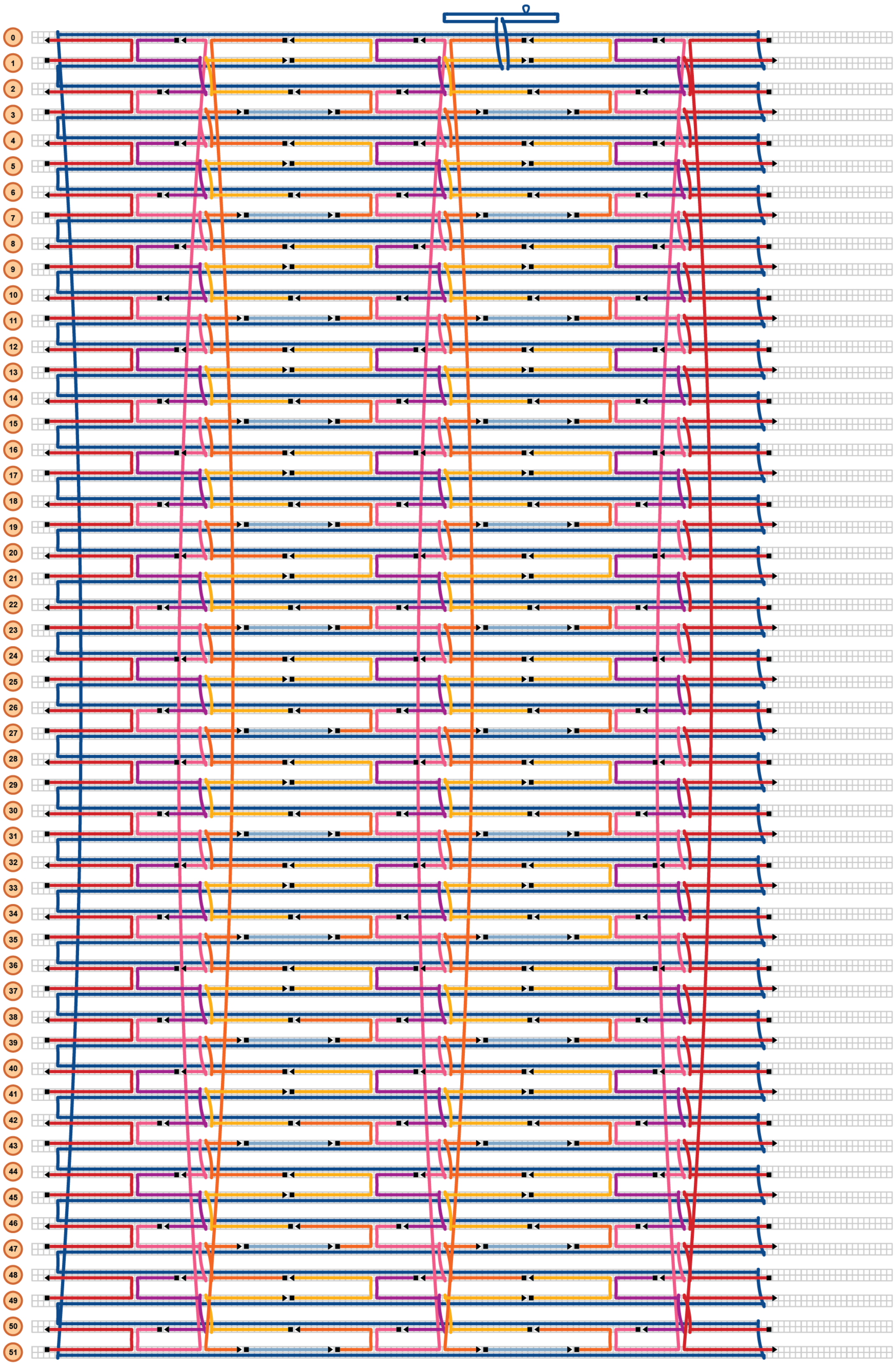


**Figure S17. Cadnano schematic of the 52-helix pleated nanotube.**

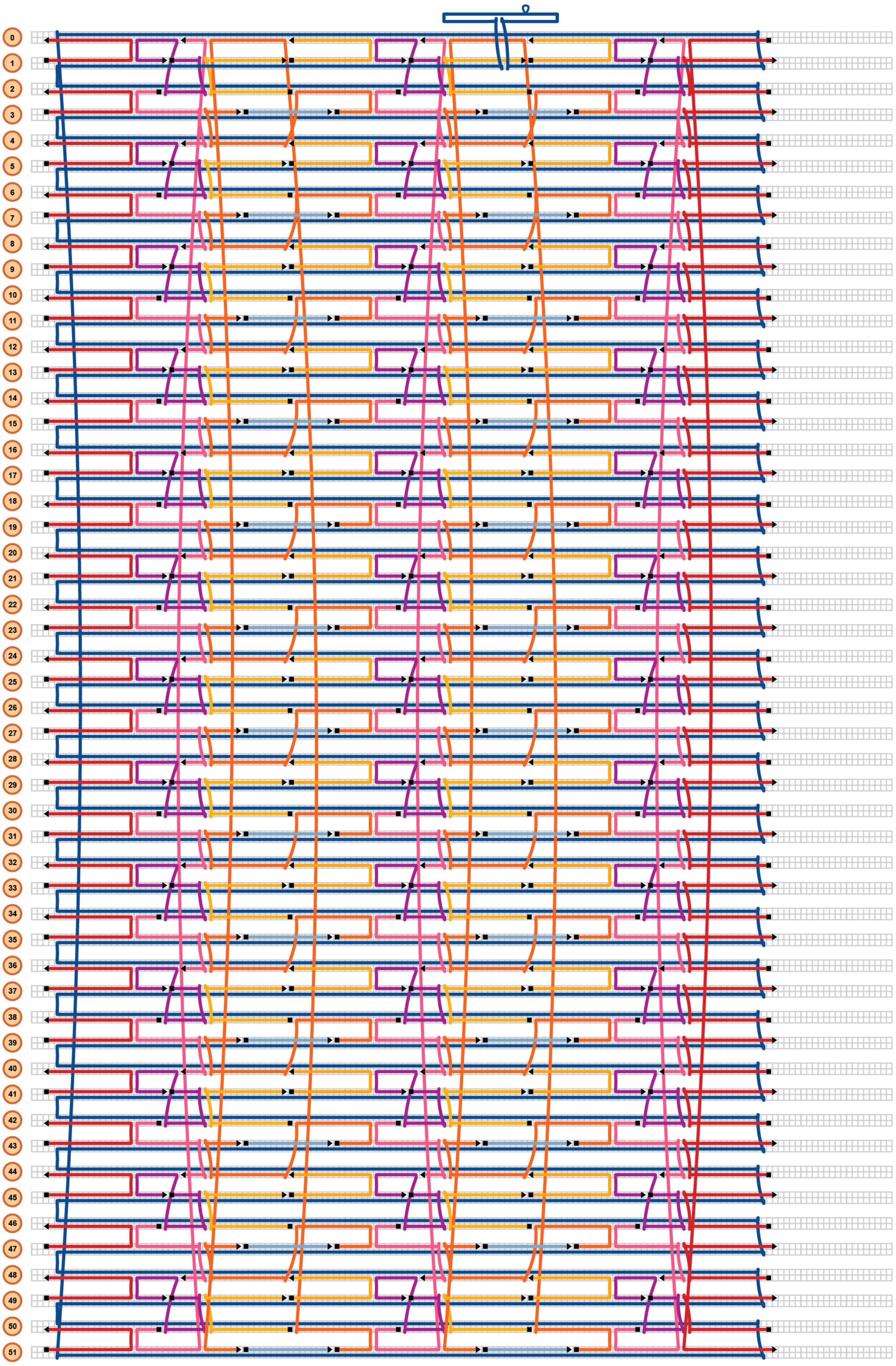


**Figure S18. Cadnano schematic of the 52-helix pleated APA-stapled nanotube.**

**Table S1. Summary of predicted and measured nanotube geometry.**

| Nanotube Design | Measurement Type | Predicted | TEM Measurement | AFM Measurement | Cryo-EM* Measurement |
|---|---|---|---|---|---|
| 10-helix pleated | Length (nm) | 237.7 | 230.2 ± 2.5 | | |
| | Diameter (nm) | 11.3 | 8.5 ± 1.1 | | |
| 20-helix | Length (nm) | 119.7 | 118 ± 2 | 120.1 ± 4.5 | 118.1 ± 3.4 |
| | Diameter (nm) | 21.2 | 20.6 ± 1.1 | 30.4 ± 2.6 | 19.4 ± 1.7 |
| 36-helix pleated | Length (nm) | 65.3 | 63.5 ± 1.6 | 60.3 ± 3.6 | 63.1 ± 2.7 |
| | Diameter (nm) | 24.2 | 28 ± 1.6 | 36.8 ± 5 | 25.2 ± 2.8 |
| | Circumference (nm) | 76.1 | | | 81 ± 1.6 |
| 36-helix pleated APA-stapled | Length (nm) | 65.3 | 63.4 ± 2.2 | 57.8 ± 3.3 | 60.2 ± 4.7 |
| | Diameter (nm) | 24.2 | 24 ± 1.6 | 28.9 ± 3.7 | 23.2 ± 2.3 |
| | Circumference (nm) | 76.1 | | | |
| 36-helix pleated APA1t-stapled | Length (nm) | 65.3 | 63.2 ± 2.7 | 58.9 ± 4.1 | 64 ± 4.1 |
| | Diameter (nm) | 24.2 | 25.1 ± 2.1 | 29.2 ± 5.2 | 21 ± 3.2 |
| | Circumference (nm) | 76.1 | | | 73.7 ± 5.5 |
| 52-helix pleated | Length (nm) | 42.3 | 43.1 ± 1.4 | 35 ± 7.6 | 41.7 ± 2.2 |
| | Diameter (nm) | 31.9 | 43 ± 2.6 | 57.1 ± 5.2 | 39.8 ± 3.4 |
| | Circumference (nm) | 100.2 | | | 115.5 ± 4.6 |
| 52-helix pleated APA-stapled | Length (nm) | 42.3 | 43.1 ± 1.3 | | 43 ± 2.9 |
| | Diameter (nm) | 31.9 | 34.9 ± 1.5 | | 33 ± 6.1 |
| | Circumference (nm) | 100.2 | | | |
| 52-helix pleated APA1t-stapled | Length (nm) | 42.3 | 42.4 ± 1.2 | 31.7 ± 3 | 42.2 ± 2.1 |
| | Diameter (nm) | 31.9 | 37.1 ± 1.9 | 42.2 ± 3.6 | 32.8 ± 4.1 |
| | Circumference (nm) | 100.2 | | | 104.7 ± 4.4 |

*Data for Cryo-EM measurements of pleated nanotubes with APA1t staples are shown in Supplementary Figure S11

1. Douglas, S.M., Marblestone, A.H., Teerapittayanon, S., Vazquez, A., Church, G.M. and Shih, W.M. (2009) Rapid prototyping of 3D DNA-origami shapes with caDNAno. *Nucleic Acids Res.*, **37**, 5001-5006.
